# Integrative Analysis of the Mouse Cecal Microbiome Across Diet, Age, and Weight in the Diverse BXD Population

**DOI:** 10.1101/2025.07.31.667922

**Authors:** Ziyun Zhou, Arianna Lamanna, Rashi Halder, Emeline Pansart, Shaman Narayanasamy, Besma Boussoufa, Thamila Kerkour, Paul Wilmes, Evan Williams

## Abstract

**Background:** The gut microbiota adapts to and shapes the host’s metabolic state through affecting circulating metabolites and consequent gene regulatory networks, resulting in systemic influences in diverse organs via connections such as the gut–liver axis. Numerous variables such as diet, age, and host genetics modulate the composition of the gut microbiome, but their interactions and specific associative and mechanistic links to host molecular phenotypes remain incompletely unannotated. Integrated multi-omics approaches in genetically diverse populations offer an opportunity to dissect these interactions and identify predictive microbial signatures for host phenotypes, such as body weight, and molecular associations with gene expression pathways in gut and liver.

**Results:** We sequenced, aligned, and integrated the cecal metagenome, metatranscriptome, and host transcriptome from 232 mice across 175 distinct cohorts according to a low fat chow diet (CD) or a high-fat diet (HF), four adult ages (between roughly 180 to 730 days of age), and 43 distinct genotypes (inbred BXD strains). Genetics and diet exerted the strongest influence on microbiota abundance and activity, followed by age. HF feeding significantly reduced diversity across all ages and all genotypes, altering >300 species. Machine learning models based on microbial profiles reliably predicted body weight within dietary group (AUC = 0.84 for CD, 0.79 for HF) and chronological age (AUC = 0.84), with model performance rising of age prediction rising to 0.95 when integrating top microbial features with liver proteomics. Network analyses of expression data revealed links between genes, pathways, and specific microbes, including a negative association between cecal *Ido1* expression and short-chain fatty acid (SCFA)-producing Lachnospiraceae, suggesting dietary fat may modulate host tryptophan metabolism through microbiota shifts.

**Conclusions:** Whole metagenome and metatranscriptome sequencing approaches have massively expanded the landscape of microbiome analysis compared to earlier short-read 16S analyses. The resulting datasets quantify hundreds of uniquely identifiable microbes, which can be used to create sets of highly predictive microbial biomarkers for aging and obesity. When trained on controlled mouse populations, these results demonstrate that microbiome profiling can achieve high predictive capacity (AUC = 0.95 with multi-omics integration) for complex readouts such as age and body weight (AUC = 0.84), even considering genetic and dietary variation, establishing a framework for biomarker development. While at present many bacteria are still functionally unannotated at the species level, multi-omics approaches — including gene expression from the host tissues — provide insights into the functional associations of specific taxa in the microbiome.

## Introduction

The typical, individual mammalian intestinal tract is colonized by trillions of bacteria, coming from thousands to tens of thousands of species, collectively called the gut microbiota [1]. Acting as a crucial link between diet and host metabolism, gut microbiota play an essential role in digestion and nutrient absorption and can impact overall metabolic health [2]. For instance, “germ-free” mice born and raised in sterile environments do not become obese when fed a high-fat and high-carbohydrate diet [3], but they also have immunodeficiencies [4], and with age they tend to suffer from a variety of metabolic disorders such as a dramatic hypertrophy of the cecum [5]. In parallel with diet, aging also alters the diversity and stability of gut microbiota [6]. These aging-associated changes are linked to increased inflammation and a reduction in beneficial metabolite production, which may further accelerate metabolic and immune decline [7].

The cecum, located at the junction between the small intestine and the colon, is a notable region for microbial activity, as it serves as a primary fermentation site for indigestible fibers, particularly for the production of short-chain fatty acids (SCFAs) such as butyrate [8]. The cecum is anatomically highly distinct, facilitating equivalent sample collection over long study periods (6 years in the case of our colony). The cecum is a “cul-de-sac” and has a relatively long residence time, with it filling by 6 hours after feeding and then emptying over the next 16 hours [9]. The relative ease of precisely-consistent tissue collection makes the cecum an ideal source for microbial sampling for long-duration experiments. Furthermore, and in contrast to collecting fresh feces from the rectum, cecal collection permits measurement of both the microbe content and the gene expression networks of immediately-surrounding mouse tissue, which can offer direct insights into systemic host-microbiota interactions that are essential for understanding the etiology of metabolic diseases [8, 10]. Conversely, cecal tissue in mice cannot be taken longitudinally, necessitating the use of inbred strains to examine how the cecal microbiome changes across age, diet, or other interventions.

Numerous clinical studies over the past decade have shown that the gut microbiome as a whole can have a causal effect on both causing and curing complex metabolic diseases. For instance, mice with inflammatory bowel disease (IBD) can be cured by oral delivery of specific *Akkermansia* bacteria [11], and fecal microbiota transplants (FMT) have been widely proven to cure *Clostridioides difficile* infections in humans, which hitherto had been notoriously difficult to treat [12]. However, given the high diversity of the microbiome and the colonial nature of microbe-microbe interactions, research in this field has often struggled to identify reliable causal mechanistic relationships between particular microbial taxa to predictable phenotypic outcomes. Successful clinical studies using FMT result in whole-scale shifts affecting hundreds of billions of bacteria, meaning that the specific causal bacteria and their mechanisms are exceedingly difficult to identify, validate, standardize, and reproduce at scale. That specific taxonomic shifts are relevant to diagnosing and treating diseases related to the digestive system like IBD and colon cancer are perhaps unsurprising, yet in recent years, numerous pioneering clinical studies have further found the microbiome to be causally associated with a dizzying variety of diseases far more distal to intestinal health, from Parkinson’s disease [13] to type I diabetes (T1D) [14] and type II diabetes (T2D) [15]. However, performing FMT to treat such systemic diseases, and metabolic phenotypes such as morbid obesity, have had unpredictable results [16]. More detailed mechanistic understanding of the relationship between the microbiome, host, and metabolic health is thus necessary to develop new, targeted treatments.

Therefore, in this study, we generated multi-omics data in the ceca from 232 mice varying according to the interactions between our three independent experimental variables: age (between 157 to 828 days), diet (low fat “chow” (CD) or high fat (HF)), and genotype (43 inbred strains from the BXD mouse genetic reference population (GRP)). In this cohort, we measured the metagenome (MG) and metatranscriptome (MT) from the cecal contents, and the transcriptome of the cecum tissue itself (abbreviated as “mRNA”). These newly generated data are then supplemented with our prior phenotype data on these mice, and their individual liver transcriptome and proteomes [17]. Here, we aim to first show that each molecular layer of data is reliable and brings distinct analytical insight into the metabolism of these mice. Then, we seek to identify and quantify the abundance and activity of specific microbial taxa associated with diet, body weight, and age. Lastly, we apply both traditional bioinformatics approaches and newer machine learning methods to calculate the relationship of these datasets and their interactions to predict key metabolic phenotypes like obesity and expression of key gene networks, and to identify putative mechanistic pathways and functions. We show that these multi-tissue, multi-omic data can be used to predict major phenotypic parameters such as body weight, and also provide insight into the cellular activity of other tissues in the mouse. That is, not only can we identify specific associations between the microbiome and metabolic health, but we can also infer the affected mechanistic pathways and — by using the different ages of cohorts — indicate the consistency and timeline of the connection.

## Results

### Optimized Isolation of Metagenomic and Metatranscriptomic Data from Preserved Cecal Samples in a Diverse Aging Mouse Population

In 2021, we completed the phenotyping of an aging population of 2157 mice from 89 BXD strains [18], tested in two diets with calories obtained predominantly from carbohydrates (i.e. CD) or from fats (i.e. HF). 662 of these individuals were taken for tissue collection at four target intervals: 200, 360, 550, and 730 days of age (i.e. in steps of 6 months). Our laboratory has recently completed a detailed molecular analysis of the paired transcriptome, proteome, and metabolome in 347 livers of these mice from a subset of 58 strains across diet and age [17]. In that study, we were able to identify predictive molecular signals of a mouse’s future outcomes (such as lifespan), as well as regulatory genes affecting health (such as susceptibility to diet-induced obesity) and the control of molecular pathways affecting aging and diet related genes (such as cholesterol synthesis). From that aging study, we took the whole, flash-frozen cecum samples from 597 individuals in 56 different strains (all overnight fasted, CD n = 299, HF n = 298). For those 597 individuals, there are typically two biological replicates for the selected strain-diet-age matches. From this group, we selected the cecum from a subset of 232 mice from 175 cohorts (i.e. distinct strain, diet, and age groups) in order to study a wide range of interacting variables in a cost-conscious manner. For these cecal samples, we have sequenced their contents for MG (n = 200 mice, of which 10 were performed in replicate), MT (n = 80), as well as the cecal mRNA itself (n = 80) (Figure 1 A, Table S1). The same exact mice were selected for all MT and mRNA samples, while 48 overlapped between MT and MG, and overall 155 of these samples overlapped with our prior liver transcript (140 overlap) or protein expression data (138 overlap) (Figure S1 A, exact list of samples overlapping shown in Table S1 sheet 2). Dataset sizes are shown in Table 1). An overview of the experimental and analytical workflow, including extraction, sequencing, quality control, and downstream processing, is presented in Figure 1 B. After cleanup, the resulting relative proportions of the microbial to host DNA content for each sample were estimated using RT-qPCR, and samples were subsequently ranked and filtered based on these ratios (Figure S1 B-C). For RNA quality control, all samples were run on a Bioanalyzer to check for the presence of four distinct rRNA bands, indicating adequate abundance of both mouse-derived RNA for cecal mRNA measurements and microbe-derived RNA for MT measurements (Figure S1 D). The RNA samples were then separated based on poly(A) enrichment, with the poly(A)-enriched sample being sequenced to obtain the cecal tissue mRNA.

**Figure 1.**
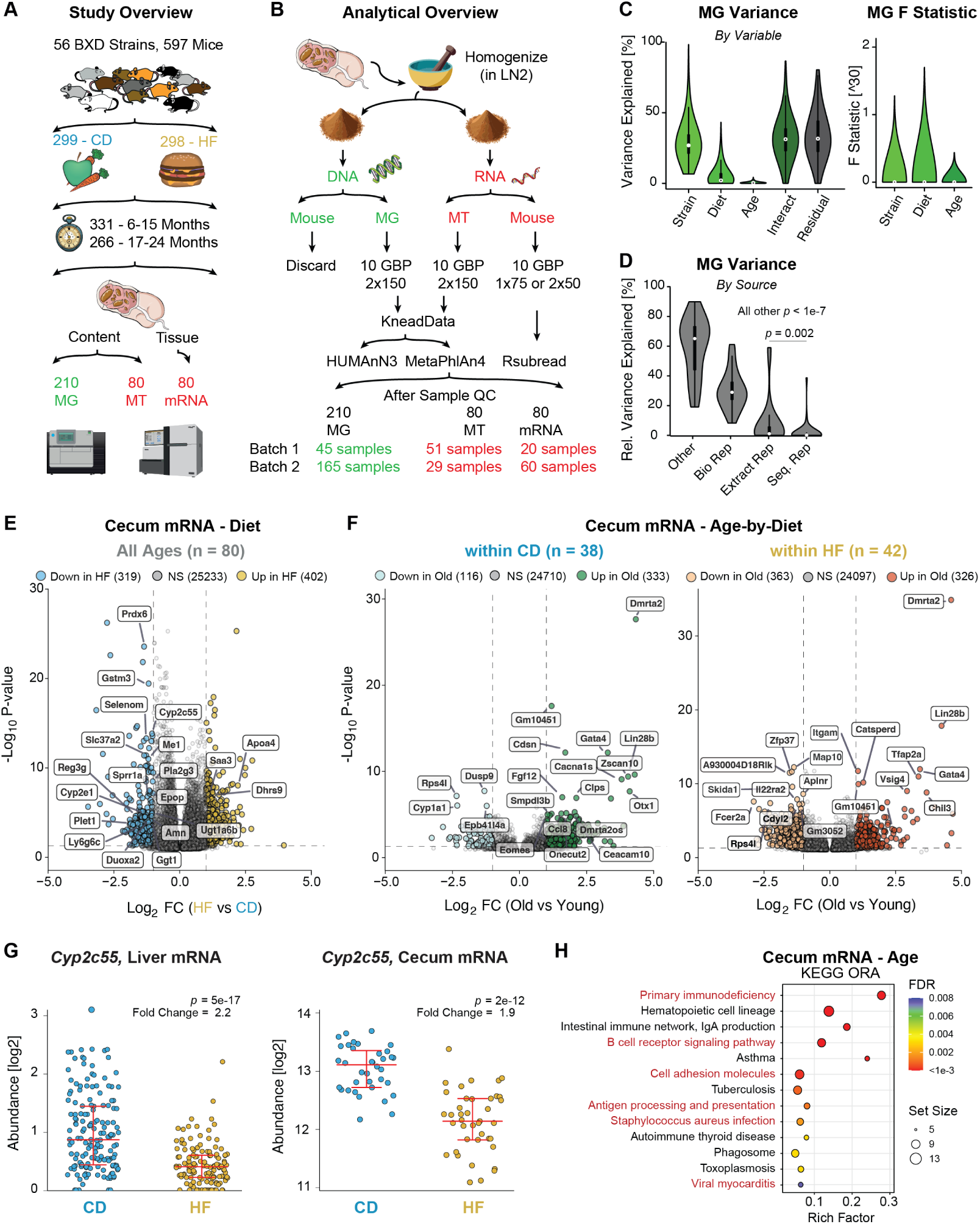
Experimental workflow and analysis of multifactorial cecal datasets. **(A)** Study design for mouse phenotyping and data generation. (**B**) Experimental process for extracting, separating, sequencing, and analyzing the cecal samples. GBP is “gigabase pair” for how many reads were done per sample, NxY is for single (N=1) or paired (N=2) end reads of length Y basepairs. (**C**) ANOVA results for variance and F statistic (adjusted for degrees of freedom) for the 34 most abundant genera in MG data. “Interact” represents the combined effects of all interactions: Strain*Diet, Strain*Age, Diet*Age, Strain*Diet*Age. (**D**) Variance explained for the same MG data comparing general variation, to that within biological replicates, to within extraction replicates, to within re-sequencing replicates. (**E**) Volcano plot showing diet-associated DEGs in cecal mRNA. Blue and yellow dots indicate genes down- or upregulated in HF (reference = CD), respectively, while grey dots represent non-significant (NS) genes. The 20 most significant genes that are also in the top quartile of expression are labeled (**F**) Volcano plots showing age-associated DEGs by diet group. Blue and yellow indicate genes downregulated with age, while green and orange represent upregulation. (**G**) Dot plot showing the diet effect on *Cyp2c55* expression in liver and cecum in the same cohort. (**H**) The 15 most significantly age-associated KEGG pathways, based on DEGs from the cecal mRNA dataset.

**Table 1.** Overview of datasets and sample counts used in this study.

| Dataset | Samples<br>(n) | Diet |  | Age (days) |  |  |  | Strain | Features | Metadata source |
| --- | --- | --- | --- | --- | --- | --- | --- | --- | --- | --- |
|  |  | CD | HF | <250 | 250-449 | 450-649 | ≥650 |  |  |  |
| MG | 200 | 112 | 88 | 62 | 46 | 47 | 45 | 43 | 614 | Table S1,<br>second sheet |
| MT | 80 | 38 | 42 | 40 | 0 | 0 | 40 | 20 | 532 |  |
| CT | 80 | 38 | 42 | 40 | 0 | 0 | 40 | 20 | 24573 |  |
| LT | 291 | 162 | 129 | 75 | 90 | 87 | 39 | 54 | 25394 | Previously<br>published<br>dataset [17] |
| LP | 316 | 165 | 151 | 78 | 105 | 82 | 51 | 57 | 3940 |  |

After read alignment, counting, and normalizing, we used ANOVA to systematically evaluate the relative contributions of the three independent variables — genotype (“strain”), diet, and age — to the observed variation in our multi-omics data, across the MG, MT, and cecal mRNA datasets. These results revealed that genotype, diet, age, and their interactions together explained, on average, 60-70% of the observed variation in MG (Figure 1 C), MT, and cecal mRNA datasets (Figure S1 E). While strain had by far the largest impact on variance, it is important to note the difference in degrees of freedom between the independent variables. The samples spanned 43 BXD strains, but only two diets, and the ages of sacrificed animals were clustered specifically around four timepoints in adulthood. To address this imbalance in the degrees of freedom for each class of independent variable, we calculated the F-statistic, which is adjusted for that concern. This showed that diet individually exhibited the strongest adjusted-association with gut microbiota composition, although even once adjusted by degrees of freedom with the F-statistic, strain has almost as large of a role (Figure 1 C).

To further identify sources of variation in the MG data, 28 cohorts had biological replicates, 3 mice had identical aliquots performed in 2 extraction replicates, and 7 of the exact same DNA samples were sequenced twice on different days. While re-sequencing and re-extraction played only a very minor contribution to average measurement variation (roughly 0.5% and 2% respectively), biological variation accounted to around 30% of total variation in MG measurements (Figure 1 D). The high variability of microbiome abundances even within biological replicates, and the major effect of diet on the gut microbiome, are expected and well-documented, given that even within the same person measured longitudinally. To mitigate potential confounding effects arising from the strong diet-microbiota effect, subsequent analyses targeting diet utilized the full dataset, age-or strain- related analyses were separately performed within the CD or HF dietary groups, and for multi-omics analysis only exact mice with measurements at multi-omics measurements were considered “overlapping”. The sample sizes of each group will be indicated for each subsequent analysis.

Lastly, to evaluate the internal consistency and reliability of the gut microbiome datasets, we examined genus-level correlations between MG and MT profiles for the 48 overlapping samples. The correlation distribution was strongly shifted toward positive values compared to a shuffled control (median correlation of around 0.75), indicating strong concordance between DNA- and RNA- based taxonomic estimates (Figure S1 F). In addition, principal component analysis (PCA) of species-level MG profiles reliably overlap between the two chronological extraction batches, indicating that batch control was effective (Figure S1 G). Similarly, both batches of cecal mRNA showed no significant differences after both were transformed and variance stabilized (Figure S1 H). Together, these results demonstrate high internal agreement and minimal batch effects, supporting the reliability of the MG, MT, and cecal mRNA datasets for downstream integrative analyses.

### Diet and Aging Alter the Cecal mRNA

Host cecal mRNA data were aligned and quantified using `Rsubread` [19], and normalized gene counts were used for downstream analysis (Figure 1 B, Tables S2 and S3). Here, we aim to connect gene expression mechanisms to bacterial abundances to pathway activities and phenotypes, particularly body weight. To do so, we first analyzed the direct effect of our key independent variables of diet and age on mRNA expression in the ceca (Figure 1 E-F). Differential expression analysis of cecal mRNA revealed distinct transcriptional responses to both diet and age. A total of 721 significant differentially expressed genes (DEGs) were observed (FDR < 0.05, |log_2_FC| > 1) between CD and HF groups, including 319 downregulated and 402 upregulated by HF, Figure 1 E). To assess data reliability, we examined highly-expressed genes (top quartile) that were also the most significant and with the largest fold changes. This category of genes included many notable genes affected by diet, such as *Prdx6* [20], *Selenom* [21], and *Cyp2c55* [22] were among the most strongly reduced by HF feeding, while *Apoa4* [23] and *Saa3* [24] were upregulated.

We next examined age-associated transcriptional changes, separately for each dietary condition. Within CD cohorts, 449 DEGs were linked to age (Figure 1 F, left panel), while 689 DEGs were linked with HF cohorts (Figure 1 F, right). In contrast to diet, the most-abundant and most-significant genes in the age analysis were less consistently reported in literature, with only limited prior knowledge for genes such as the upregulation of *Ceacam10* [25] with age and the downregulation of *Lars2* [26]. To obtain a larger set of DEGs affected by age, we compared the 20 DEGs most affected by age from the CD and HF age subsets (even if not in the highest quartile of expression). Among the genes upregulated in old mice, five were shared in both diets: *Dmrta2*, *Lin28b*, *Gata4*, *Onecut2*, and *Gm10451*. In contrast, only *Rps4l* and *Lars2* were consistently upregulated in young mice under both diets. In addition to *Lars2*, two more of these overlapping genes are known to change with age: *Gata4* [27] and *Rps4l* [28]. Consistent links with age independently of diet and across 20 distinct BXD genotypes suggests an underlying cellular relationship, even in the absence of literature. Given that diet has been far more extensively studied in mouse studies of gene expression, we considered the strong overlap with expectation in the dietary analysis indicative of data reliability of age-related DEGs. While certain gene pathways may be specific to the cecum, we have also recently measured the liver transcriptome and proteome in the same mice [17], allowing us to further check for cross-tissue consistency for DEGs, e.g. that *Cyp2c55* mRNA expression lowers with HF in both the liver (fold change = 1.4) and cecum (fold change = 1.9) (Figure 1 G).

To further explore functional pathways associated with age and diet as a final sanity check before integration with the microbiome data, we applied over-representation analysis (ORA) based on DEGs and gene set enrichment analysis (GSEA) using ranked differential expression values on the cecal data. For diet-associated changes (FDR < 0.05), ORA identified 24 significantly enriched pathways, while GSEA revealed 99 pathways. 16 pathways were consistently detected by both approaches. The top 15 enriched pathways from each method are shown in Figure S2 A–B, with overlapping pathways highlighted in red. In addition, KEGG pathway analysis revealed that several metabolic pathways previously linked to HF diet in literature were significantly affected, including less intuitive pathways such as the “neuroactive ligand – receptor interaction” pathway [29], and more typical HF-related pathways such as “metabolism of xenobiotics by cytochrome P450”, “retinol metabolism”, and “steroid hormone biosynthesis” [30]. Decreased expression of the steroid hormone pathway may stem from HF-induced mitochondrial dysfunction that impairs cholesterol transport into mitochondria, the first step of steroid hormone biosynthesis [31]. In particular, this pathway was enriched with 15 DEGs (e.g., *Cyp2c55*, *Cyp3a11*, *Cyp1a1*, *Cyp2e1*, *Ugt1a9*) (Figure S2 C, upper panel). Additionally, the “neuroactive ligand–receptor interaction” pathway has been further reported to be upregulated by HF in the hypothalamic transcriptome of mice [32], potentially driven by the diet–gut–brain axis. In our data, this pathway was highly enriched (NES = 2.05, p = 2e-9) with key DEGs including *Lep*, *Grik1*, *Avpr2*, and *Grm7*, among others (Figure S2 C, lower panel).

Building on the same analytical framework, we next investigated how age influences pathway-level changes through both ORA and GSEA (Figure 1 H, Figure S2 D). ORA identified 27 significantly enriched pathways, while GSEA revealed 58, with 19 pathways consistently detected by both methods. Among these, the B cell receptor signaling pathway was significantly enriched in both analyses. This finding aligns with previous reports in humans and animal models, which have shown that aging is often accompanied by a reduction in B cell numbers and functional impairments within B cell populations [33, 34], with 10 DEGs according to age in the cecal samples (Figure S2 E, left panel). Conversely, several significant pathways positively associated with aging have also been extensively studied, such as the NOD-like receptor signaling pathway (Figure S2 E, right panel), which is upregulated with age but can be effectively suppressed by fecal microbiota transplantation from young donors [35]. These cecal mRNA results reveal distinct pathway alterations driven by diet and age, offering a potential framework for integration with microbial functional data.

### MG and MT Analyses Reveal Cecal Microbiome Changes Associated with Dietary Change, Aging and Strains

We next examined the broad taxonomic profiles of the gut microbiota, using MG and MT reads processed with KneadData [36] and analyzed using MetaPhlAn4 [37] (Figure 1 B, Tables S4 and S5). HF diet reduced the variety of identified taxa by around 30 to 50% across all taxonomic levels in both MG and MT data (Figure S3 A). In addition, overall Shannon α-diversity was consistently around 20% lower in HF-fed mice across all age groups, with a comparable reduction observed in both MG and MT datasets (Figure 2 A). These findings are consistent with previous widely reported observations that a HF diet reduces microbiome diversity rapidly – typically in under a month – which includes the time on HF for the earliest timepoint in our study (roughly 90 days after the start of HF feeding) [38]. Overall, 39% of observed variation in α-diversity was due to diet, 20% due to genotype, and perhaps 1% due to age (by ANOVA; p < 2e-16 for diet, p = 1e-6 for genotype, and a suggestive p = 0.06 for age). Notably, age had no meaningful effect: α-diversity reduced by around 20% at all ages, despite the lengthened time on HF. Conversely, β-diversity analyses (Bray-Curtis dissimilarity) demonstrated that both diet and age significantly influenced gut microbial community structure (Figure S3 B). While diet explained a larger proportion of the variance in community composition and was associated with more pronounced shifts across samples, age accounted for a smaller portion of the variance but still exhibited statistically significant effects.

**Figure 2.**
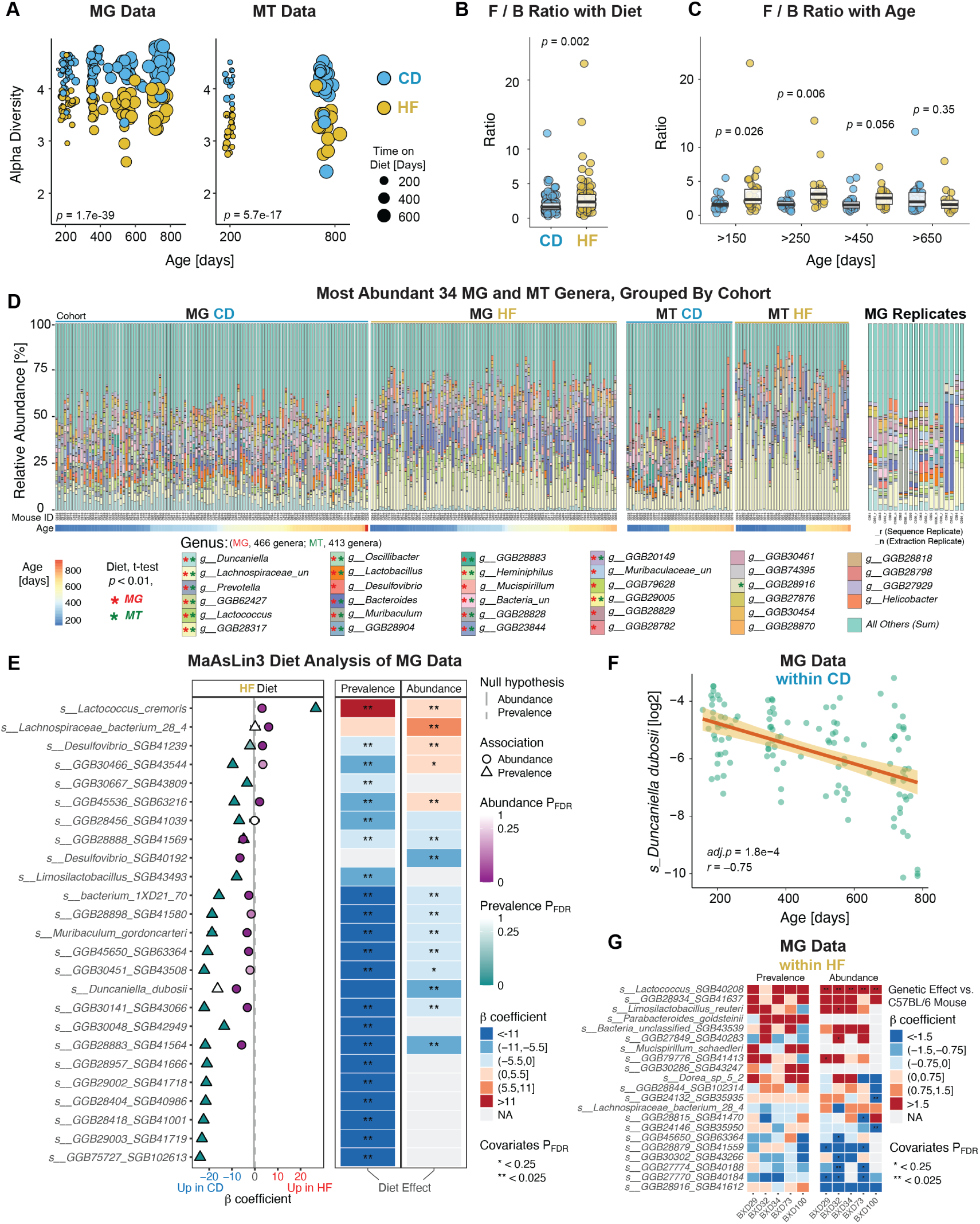
Microbial Diversity and Composition Are Shaped by Diet, Age, and Genotype in MG and MT Datasets. **(A)** The Shannon index α-diversity at the MG and MT levels, showing significantly lower diversity in HF. *P*-values indicate overall differences between diet groups across all ages. (**B**) The F/B ratio is significantly increased in HF cohorts. (**C**) The same data as panel B, now separated by age. (**D**) Stacked bar plots of the relative abundance of the 34 most abundant genera in the MG (left panels) and MT (right panels) datasets, ranked by diet-associated t-test significance. Stars indicate genera with adjusted *p*-values < 0.01. MG samples performed as technical replicates are at the far right. (**E**) The top 25 species most significantly associated with diet in the MG dataset, identified by MaAsLin3. Direction and significance of associations are shown for prevalence and abundance. (**F**) *Duncaniella dubosii*, the most strongly age-correlated taxon identified by MaAsLin3, shows a significant negative correlation with age. (**G**) Correlation matrix showing strain-level differences in species abundance, using B6 as the reference.

We then checked whether taxonomic signatures in the ceca will reliably indicate general health outcomes, rather than only recapitulate the known dietary state of the mice. To initially test this, we checked the phyla abundance ratio between Firmicutes and Bacteroidetes (F/B), which in mice are consistently reported to track with obesity as well as HF diets [39] (although results in humans are much less consistent [40]). Here, we again observed a highly significant elevation in HF cohorts (Figure 2 B), and Firmicutes and Bacteroidetes abundances also significantly correlated with body weight (Figure S3 C). As with α-diversity, age suggestively explained only 2% of variation (p = 0.09, ANOVA) in contrast to diet (8% variance explained) and particularly strain (32% variance explained). However, age influenced the dietary effect: no significant difference in the F/B ratio was observed between dietary groups in older mice (Figure 2 C). Similar to the observation that some CD individuals had low α-diversity and some HF animals had high α-diversity, a few strains also had low F/B ratios despite being obese and vice-versa (e.g. HF-fed BXD70s are an obese 52g but have a F/B ratio of 1.2, while CD-fed BXD69s are a thin 22g but have a F/B ratio of 3.4). The prominent role of genetic background in explaining MG abundances (i.e. Figure 1 C), diversity, and the F/B ratio may – together with age diet – explain some of the inconsistencies found in meta-analyses of the F/B ratio with obesity, even in well-controlled mouse studies [41]. Moreover, both Firmicutes and Bacteroidetes are large and highly-diverse phyla of bacteria, and their lower taxonomic branches can have diverse effects. For instance, within the Firmicutes phylum, *Lactobacillus reuteri* has been associated with increased obesity, whereas *Lactobacillus gasseri SBT2055* exhibits anti-obesogenic properties [42]. These confounding factors (among others) likely explain the limited translational success of phyla-based microbiome analyses, such as the F/B ratio [40]. However, by generating MG data as in this study (or using long-read 16S sequencing), it is possible to perform more granular analysis at more specific taxonomic rankings: particularly genus, species, and potentially strain.

Before searching for novel associations with metabolic health at lower taxonomic ranks, we performed a final quality control to check for reliable, reproducible, known associations with host health. To this end, we analyzed genus-level relative abundance profiles derived from both MG and MT datasets (Figure 2 D) to dissect microbial composition. Samples were grouped by diet, and within each dietary group, individuals were ordered from left to right by increasing age. We visualized genera by taking the union of the top 20 most abundant genera from the MG and MT datasets and the top 25 genera shared between them, resulting in a total of 34 unique genera (plus “Others”, which is the sum of all smaller genera). Here, we again observe marked differences in genus composition between the CD and HF groups, such as a large decrease in *Duncaniella* in HF, and a substantial increase in a large cluster of unclassified genera within the Lachnospiraceae family (bottom two bars in light blue and pale yellow, respectively, Figure 2 D). Overall, 24 taxa were significantly regulated by diet in the MG data and 19 in the MT data, of which 18 were significant in both datasets. Conversely, only one genus varied with age in both datasets (*GGB62427*) while *Duncaniella* also varied by age in MG data, although only within CD cohorts, and *GGB28904* varied by age only within the MT data.

Besides the overall decrease in diversity with HF, specifically-affected taxonomic classifications also broadly matched expectations from literature. For instance, *Lachnospiraceae_unclassified*, *Lactobacillus*, and *Bacteroides* exhibited increased relative abundance due to HF [43, 44], while *Lactococcus* [45], *Prevotella*, and *Lactobacillus* were less abundant in HF. However, certain highly-consistent and variable genera, such as *GGB62427*, were less well-characterized and have not been extensively studied in the context of diet or age. Together, these results provide a valuable resource for identifying candidate genera consistently associated with obesity-promoting diets independently of genetic background.

Given the complex influences of diet, age, and strain, we next examined species-level profiles to uncover more specific microbial associations. Based on the results of MaAsLin3 [46], we identified a total of 274 gut microbial species at the MG level that were significantly associated (FDR < 0.05) with diet, and 31 species correlated with age. The 25 most strongly and significantly diet-associated features are displayed for the MG and MT datasets in Figure 2 E and Figure S3 D, respectively. The β-coefficients from the multivariable models indicate both the direction and magnitude of association with the HF diet (positive indicates higher in HF). Since we used relative abundance data, both abundance and prevalence were considered in the analysis. Abundance examines how the level a species is expressed (similar to prior analysis), while prevalence indicates how frequently a species is found across a population – an important distinction given that even relatively abundant species such as *D. dubosii* can be up to 10% abundance in certain mice, while it is completely undetected in 27% of the samples. Such discrepancies also often occur in cases of competitive metabolic niches: multiple bacteria may be capable of filling a particular niche, but stochastic effects or different initial starting abundances can cause differences in propagation between two individuals. Thus, of the top 25 species visualized, 12 were commonly altered in both abundance and prevalence models.

While the results often overlap, discrepancies and alignments help provide a more comprehensive interpretation of the relationship between species and diet. For example, among the top features, *Lactococcus cremoris* is not detected in 60% of samples, and it has both increased abundance and prevalence in the HF group, suggesting not only a higher overall abundance but also a broader presence across individuals in the cohort. In comparison,

*Desulfovibrio SGB41239* exhibits increased abundance but decreased prevalence under HF, indicating that while this taxon becomes more abundant in certain individuals, its occurrence became more restricted across the cohort overall. This discrepancy may reflect ecological heterogeneity within the HF group, for instance *Desulfovibrio SGB41239* had a negative prevalence change across the entire cohort, but it had an increased abundance in a subset of mice which may have distinct gut environments or host responses. In contrast, species such as *Muribaculum gordoncarteri* and *Duncaniella dubosii* were consistently depleted in both abundance and prevalence by HF, highlighting their general reduction in both quantity and distribution. The observed species-level shifts mirrored those at the genus or family level, but provided more granular identification of diet-associated gut microbes. Notably, four species appeared among the top 25 diet-associated taxa in both the MG and MT datasets (Figure S3 D). For example, the abundance of *Duncaniella dubosii* was significantly reduced by the HF in both the MG and MT datasets (Figure S3 E). Their consistent detection across MG and MT levels suggests that these taxa are not only present, but also transcriptionally active in response to dietary changes. In addition to well-characterized species, several unclassified taxa showed concordant patterns between MG and MT. For example, *bacterium_1XD21_70* consistently decreased in both abundance and prevalence under the HF diet (Figure S3 F-G), despite lacking functional annotation. These shared and reproducible species may serve as robust candidates for hypothesis generation and future functional validation.

To disentangle the effects of age and strain from dietary influences (Figure 1 C), we performed separate species-level analyses using only the CD samples from the MG dataset. Using the same MaAsLin3 method, we identified several species that were significantly associated with age, such as *Duncaniella dubosii*, which not only decreased under HF but also showed a marked decline with age (Figure 2 F). Additionally, to investigate strain-specific microbial patterns, we used C57BL/6 (B6) mice as the control group and included data from the 5 BXD strains with more than 10 measured individuals. Several species showed notable variation across different strains, highlighting genotype-specific microbial patterns that may reflect underlying host genetic influences on gut microbiota composition (Figure 2 G). Notably, *GGB27849 SGB40283* exhibited higher prevalence and abundance in BXD32, while *GGB28815 SGB41470* was more enriched in BXD100 compared to other strains. However, the comparatively high variability between biological replicates (i.e. Figure 1 D), large number of variables, and few replicates per strain means that care would be necessary before definitively declaring a strain, e.g. BXD100, as definitively being more conducive to growth of species like *Lactococcus SGB40208* compared to B6.

We next focused on functional profiling with HUMAnN3 [36] using the same MG and MT datasets at the genus level to quantify microbial pathway abundance and coverage across all samples (Tables S6 and S7). Prior studies have shown that MT-derived data, which capture microbial activities, are often more informative for linking microbial functions to host phenotypes than MG-derived data, which reflect present microbial abundances and their functional genetic *potentials* [47]. Nevertheless, MG profiling serves as a valuable reference to account for potential biases introduced during MT data generation and interpretation. A total of 447 MetaCyc pathways were detected in the MG and 391 in the MT dataset. Among the 30 most abundant pathways across both datasets, 17 pathways in MG and 11 in MT showed significant differences between dietary groups (FDR corrected t-test, adj.p < 0.01), with 8 pathways overlapping significantly between both datasets (Figure 3 A). By age, 5 pathways varied with age in the MG data and 6 with age in the MT data, 2 of which were associated with age in both layers (superpathway of L-aspartate and L-asparagine biosynthesis, and homolactic fermentation).

**Figure 3.**
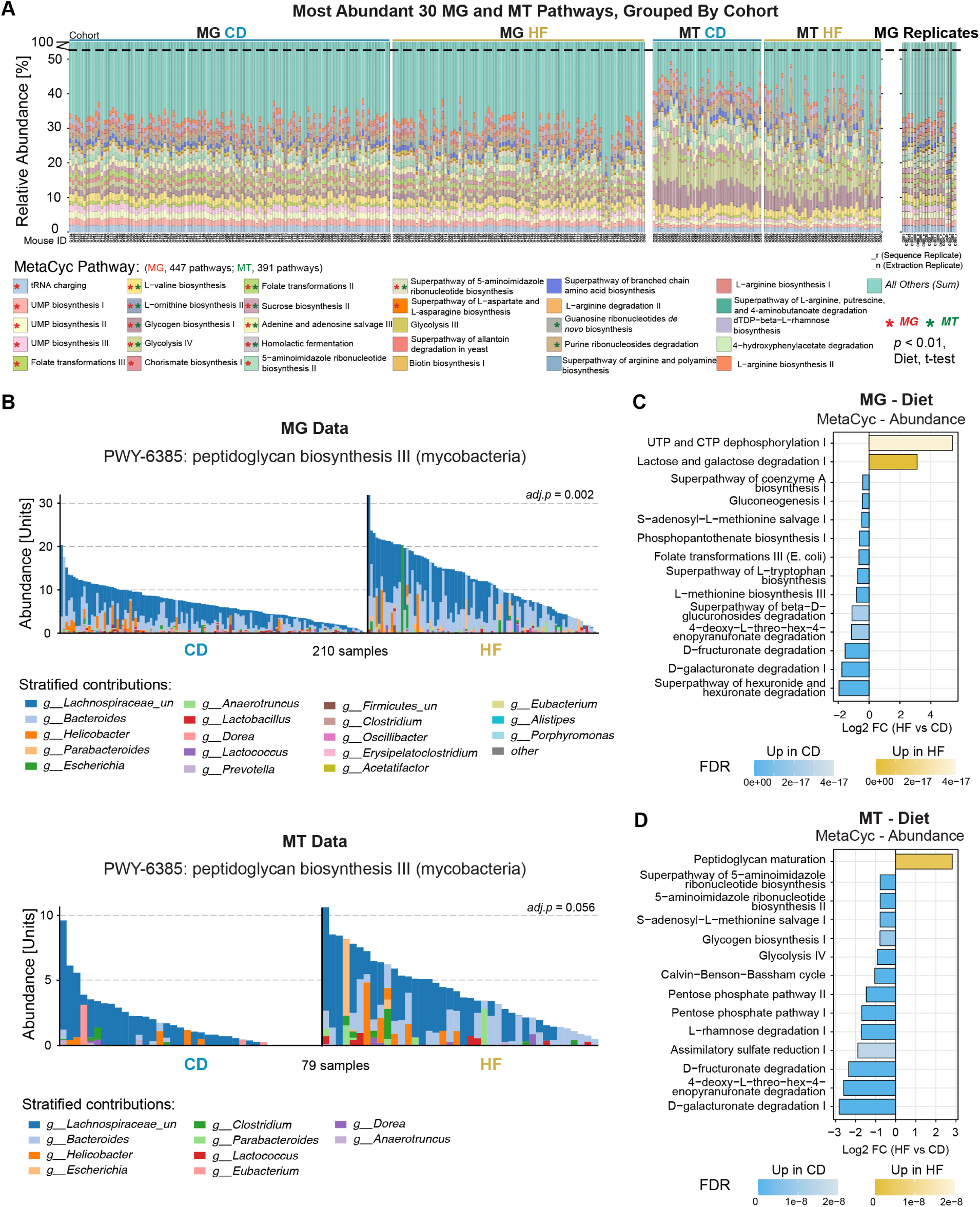
Diet-associated Changes in Microbial Functional Profiles. **(A)** Stacked bar plots of relative abundance of the 30 most abundant MetaCyc pathways in the MG (left) and MT (right) datasets, ranked by diet-associated t-test significance. Technical replicates are shown at the far right. Stars indicate pathways with adj.*p*-values < 0.01. (**B**) Dietary differences in the peptidoglycan biosynthesis III pathway based on MG (top) and MT (bottom) data. Contributions from “unclassified” bacterial genera were excluded from both panels. (**C-D**) Top 15 MetaCyc pathways with significant dietary differences identified by MaAsLin3. Abundance from MG (**C**) and MT (**D**) between CD and HF diets are shown. Bars indicate log_2_FC (HF vs CD).

In addition to these dominant pathways, several low-abundance pathways also exhibited significant associations with either diet or age, highlighting subtle but biologically relevant functional shifts. For example, the peptidoglycan biosynthesis III (mycobacteria) pathway, associated with bacterial cell wall formation and host immune stimulation, was upregulated in the HF group at both the MG and MT levels (Figure 3 B). The components of this pathway are generated by a broad range of gut bacterial genera, including *Bacteroides*, *Clostridium*, *Lachnospiraceae unclassified*, *Helicobacter*, and *Lactococcus*. In both datasets, the HF group exhibited higher total abundance of this pathway, indicating a potential upregulation of bacterial cell wall synthesis.

To comprehensively assess the associations between all pathways and host variables such as diet and age, we further performed detailed statistical comparisons using MaAsLin3, analyzing pathway abundance in MG and MT datasets (Figure 3 C-D) and pathway prevalence in the corresponding datasets (Figure S4 A-B) across different dietary groups. That is, rather than comparing the abundance and prevalence of bacterial species according to diet (i.e. as in Figure 2 E), the same analysis is performed but using functional pathways instead. The pathway abundance analysis identified 210 diet-associated pathways in MG and 110 in MT, while the pathway prevalence analysis revealed 78 significant pathways in MG and 39 in MT.

Of these, 91 pathways were shared between MG and MT in the abundance results, and 15 were shared in the prevalence results. Some pathways were primarily driven by specific species, such as “lactose and galactose degradation I” (Figure S4 C), whereas others, such as “4-deoxy-L-threo-hex-4-enopyranuronate degradation” (Figure S4 D), appeared to be largely unattributable to known species, making mechanistic investigation inherently more challenging. Notably, in both MG and MT, the “lactose and galactose degradation I” pathway was significantly upregulated in the HF, primarily driven by the marked increase of *Lactococcus lactis* under HF conditions. However, the directionality of this connection may not be universal: other studies have reported a downregulation of this pathway in HF-fed mice, which was largely attributable to a significant decrease in *Lactobacillus murinus* [48, 49]. Supporting the potential mechanistic role of this bacteria in metabolic health, a recent study supplementing mice on HF with *Lactococcus lactis* has been shown to lessen obesity and generally improve metabolic health [50]. Given that the species driving a pathway may also contribute to multiple functional modules, these results should be interpreted cautiously and are best considered hypothesis-generating, particularly to identify the mechanism and design translational studies. By contrast, the downregulation of the “4-deoxy-L-threo-hex-4-enopyranuronate degradation” pathway under HF conditions was consistent with previous research [51]. In addition, several pathways demonstrated robust results across both MG and MT (e.g., “superpathway of (R,R)-butanediol biosynthesis” and “UTP and CTP dephosphorylation I”), although no related studies have been reported to date. Together, these results provide complementary insights into microbial functional alterations associated with dietary change.

While diet plays a dominant role in all MG and MT analyses, age also significantly influenced microbial functional profiles (Figure S5 A-B). In the pathway abundance analysis (Figure S5 A), the “L-histidine biosynthesis” and “L-lysine biosynthesis III” pathways were negatively associated with aging, consistent with observations from studies across different human age groups [52]. Conversely, the “fucose and rhamnose degradation” superpathway showed a significant positive association with aging in the pathway prevalence analysis (Figure S5 B), aligning with previous findings of its upregulation in older adults with sarcopenia [53]. Taken together, these results highlight how diet and age jointly modulate the gut microbial landscape at both taxonomic and functional levels, and illustrate the utility of integrating MG and MT datasets to capture microbial metabolic outputs. While MG and MT results often align – raising confidence in the reliability of such findings – cases where they conflict can provide novel insight. Microbiome associations are also frequently distinct at the MG or MT level even when measured in the same samples [54]. In such cases, findings that are congruous at both the MG and MT level can be considered more robust, while associations that are discrepant can highlight pathways and mechanisms where microbial *abundance* does not closely track with microbial *activity*. Indeed, this may be seen as a parallel to multi-omics analyses of tissue gene expression of the transcriptome and proteome, where the majority of variation in transcript levels is not observed at the protein level [55] even though both are nominally sourced from the same gene. Discrepancies in multi-omic analyses allow unique insights into biological mechanisms, though greater care must be taken to mitigate false discovery, and follow-up validation experiments may be more difficult to design, given that they must account for why the multi-omics differ.

### Machine Learning-based Modeling to Identify Cecal Microbial Biomarkers Associated with Aging

Beyond conventional statistical analyses of taxonomic and functional profiles described in the previous section, we further employed supervised machine learning to model and predict host outcomes based on gut microbiome composition, starting with chronological age. This approach allows us to first evaluate whether microbial profiles contain sufficient predictive information about host aging, and then to identify key microbial species that contribute most to age classification. By training and validating multiple classifiers, we aimed to complement previous age-associated analyses and prioritize robust microbial biomarkers that may not be easily identified by traditional univariate methods. Given that ANOVA results (Figure 1 C) indicated a much stronger influence of diet than age on microbial composition in the MG dataset, we trained age-prediction models separately within each dietary group. Moreover, the age distribution within our cohort was not continuous (Figure S5 C), making classification more suitable than regression for downstream modeling. To ensure balanced group sizes, we defined samples younger than 350 days as the “Young” group and those older than 450 days as the “Old” group. After filtering, a total of 100 samples (Young n = 40, Old n = 60) were retained for modeling.

We trained four supervised learning models including random forest (RF), XGBoost (XGB), logistic regression (Logistic), and deep neural networks (DNN) using 5-fold cross-validation, with 80% of the samples used for training in each fold and 20% for testing, to evaluate their performance in classifying age groups from MG species-level data (614 species in total). Among these, the RF classifier achieved the highest performance with an average area under the curve (AUC) of 0.84 across folds (Figure 4 A, left panel). To further evaluate model generalizability beyond cross-validation, we conducted an independent batch-wise assessment in which one MG data batch was used for training and the other for testing. Both RF and XGB retained strong predictive performance exceeding 0.80 (Figure 4 A, right panel), supporting their robustness across biological replicates. To further enhance model expressiveness on smaller datasets, we incorporated feature interaction (FI) terms. Inclusion of interacting features yielded modest improvements in model performance (e.g. AUC-ROC improved from 0.80 to 0.88 with RF) by capturing non-linear relationships between microbial features and host age. In parallel, we also trained age classification models using MG data from the HF group and MT data from the CD group (Figure S6 A–B). In both cases, the RF classifier outperformed other models, with cross-validated AUCs of 0.81, despite the smaller sample size in the MT dataset.

**Figure 4.**
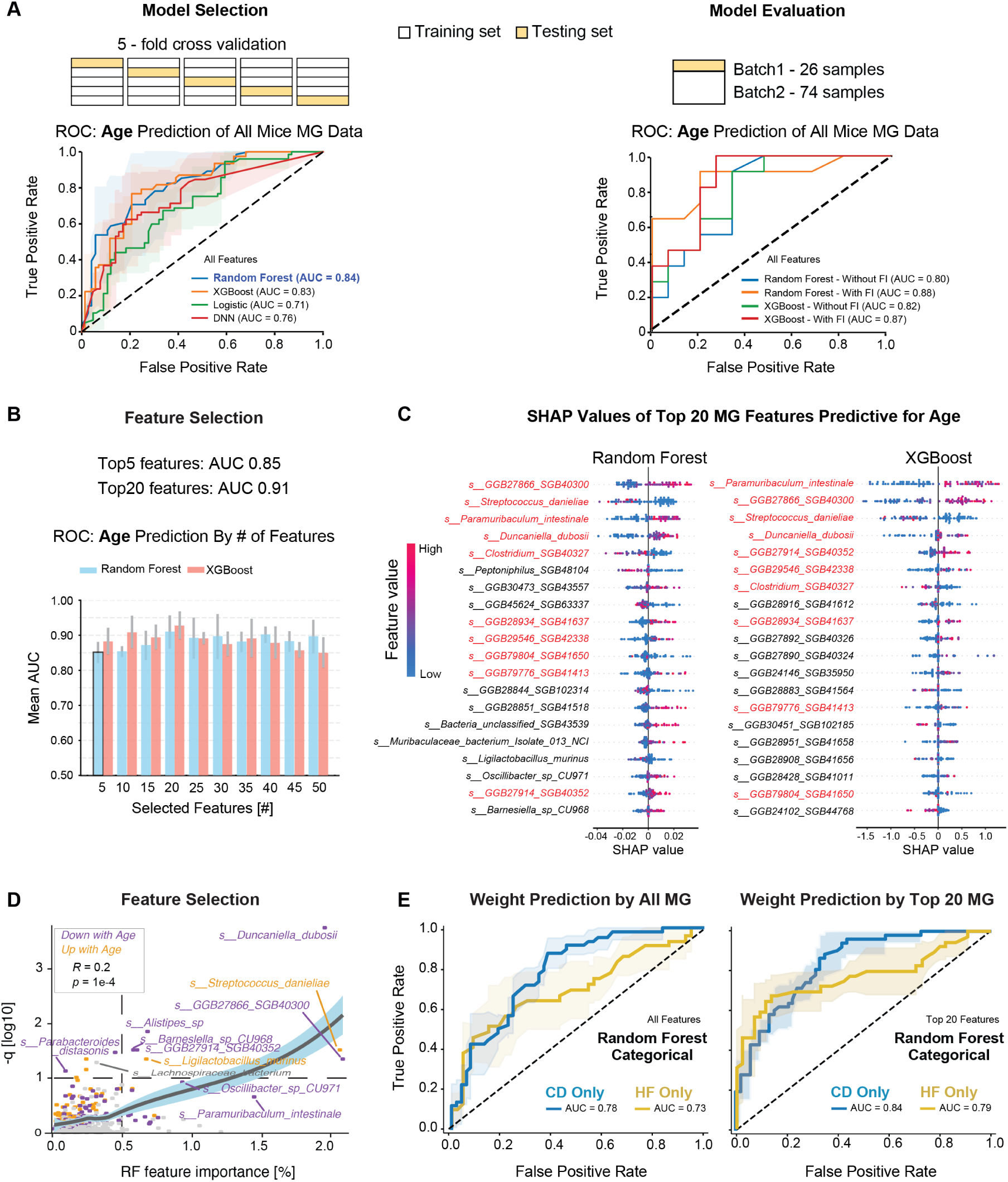
Machine Learning Models Identify Microbial Species Predictive of Host Age. **(A)** Five-fold cross-validation for model training and evaluation of age prediction using MG data across four machine learning approaches for selecting (left panel) and validation (right panel) in CD cohorts. (**B**) AUC-ROC values of models retrained using different numbers of top-ranked features with 5-fold cross-validation. (**C**) SHAP swarm plots for the top selected age-associated species, showing the direction and magnitude of each species’ contribution to age classification across samples for RF and XGB models. (**D**) Comparison of species associated with age identified by MaAsLin3 and RF. Each point represents one species. The scatter plot shows a positive correlation between RF feature importance and MaAsLin3 statistical significance. Purple points are downregulated with age, yellow are upregulated. (**E**) Results from training and evaluation of body weight prediction using RF for all features (left) or the top 20 features selected (right), analyzed within each diet separately.

Building on the robust performance of the full-feature model (AUC = 0.84), we next identified which microbial species contributed most to age classification. Feature importance was computed within the cross-validation framework and used as the basis for SHAP-derived feature ranking, with SHAP values evaluated per fold to prevent data leakage. To evaluate the contribution of top-ranked features, we retrained 5-fold cross-validation models using stepwise subsets of the top 50 ranked species, increasing the feature set in increments of five. This approach reduces overfitting risk in high-dimensional microbiome data and improves interpretability by focusing on smaller, biologically tractable sets of taxa. The RF model achieved an AUC of 0.85 using only the top 5 features, and exceeded 0.90 when using the top 20 features. No improvement was observed with larger numbers of features selected, indicating that a small subset of microbial taxa carries sufficient information for strong age prediction (Figure 4 B). XGB models showed similar trends, although their predictive performance was slightly lower than that of RF. To characterize how these top-ranked species contributed to model predictions, we visualized their SHAP values using swarm plots, which illustrate the direction and magnitude of each species’ influence across individual samples. Ranking species by SHAP value magnitudes revealed that 10 taxa overlapped among the top 20 features identified by both models (Figure 4 C), highlighting their robustness as age-associated candidates, e.g. *GGB27866 SGB40300*, *Streptococcus danieliae*, *Paramuribaculum intestinale*, and *Duncaniella dubosii*. To ensure that these selected features reflect biological aging rather than technical artifacts, such as cage- or facility-associated contamination during the extended sample collection period, we generated a combined plot of the relative abundances of the top 20 RF-ranked species across all samples. No evidence of batch-related shifts was observed (Figure S5 D), supporting their relevance to aging rather than external confounders. Overlapping features then served as prioritized candidates for generating aging-related hypotheses.

To compare microbial features identified by different analytical strategies, we next examined the overlap between statistical significance from MaAsLin3 linear models and feature importance scores from RF classifiers. While there was substantial variation between the two approaches, a subset of species was consistently highlighted by both methods (Figure 4 D). A moderate but significant positive correlation was observed between model-derived feature importance and the negative log-transformed q-values from MaAsLin3 (Spearman: *Rho* = 0.2, *p* = 1e-4), suggesting that species strongly associated with age are more likely to be prioritized across analytical frameworks. Of particular note, *Streptococcus danieliae* exhibited the strongest positive correlation with aging, whereas *Duncaniella dubosii* showed the strongest negative association. To further explore the phylogenetic distribution of age- and diet- associated microbial features, we annotated a taxonomic tree of all detected species with significance results from the MaAsLin3 analysis (Figure S6 C). This visualization was used to examine whether microbes responsive to host variables were clustered within specific taxonomic clades. However, no clear aggregation patterns were observed, suggesting that microbial responses to diet and aging are shaped by complex host–microbe interactions rather than being driven by a single functional pathway or metabolite class. Interestingly, several species highlighted by both machine learning methods and traditional bioinformatic approaches exhibited low mean abundance (e.g. *Duncaniella dubosii* is only about 1.6% of mean abundance), reinforcing the importance of using complementary analytical strategies to identify robust features. This integrative approach allowed us to uncover microbial signatures that may otherwise be overlooked – such as low-abundance bacteria with only sequential identifiers rather than formal species names (e.g. the age-associated *SGB40300*) – and offers a framework for identifying and selecting candidates for mechanistic experiments.

To evaluate the reliability and biological relevance of our modeling framework on a dependent variable (as compared to age, which was a selected independent study variable), we applied the same machine learning pipeline to predict mouse body weight using MG data (Figure 4 E). Although diet exerts a much stronger effect than age on MG dataset, and is directly associated with body weight, this strong diet signal is removed when models are trained within each diet group. As a result, body-weight prediction based on within-diet MG data shows lower performance than age prediction using the full features (Figure 4 E, left panel; RF AUC = 0.78 CD and 0.73 HF). When using feature selection to pick top 20 features, the mean AUCs improve (right panel; RF AUC = 0.84 CD and 0.79 HF), showing the utility of performing feature selection. The top 20 features ranked by RF and within each diet are shown in (Figure S6 D), with only 3 common predictive species for weight between the two diets. Note that body weight here is treated as binary classifier in order to compare with the age models, as the ages in the study cohort were centered on four discrete timepoints. For CD, 49 animals were under 25g and 55 were over 26g, and for HF 40 animals were under 39 g and 39 animals were over 45g. This result not only highlights the predictive capacity of microbiome-based features for host metabolic traits (in this case, body weight) but also indirectly supports the robustness of our age prediction models, and that the approach can be applied to both dependent (body weight) or independent (age) variables.

### Integrated Multi-omics Analyses Reveal Host-Microbiome Interactions in Metabolism and Aging

Having performed a series of both combined and independent MG and MT analyses according to the key independent variables of genotype, age, and diet in the previous sections, we identified multiple microbial changes associated with each variable. However, ultimately these taxonomic shifts should be mediated through cellular changes in the host organism before leading to phenotypic outcomes like increased body weight. To further leverage the multi-omics datasets generated from the same BXD mice [17], we conducted integrative analyses across microbiome (MG and MT), cecum (mRNA), and liver (mRNA and proteome) datasets. This enabled us to explore host–microbiome interactions more comprehensively and evaluate the added value of multi-omics integration in predicting biological traits and identifying functionally relevant microbial associations. We first examined the relationship between gut microbial composition and host metabolic gene expression by integrating MG species-level profiles with liver transcriptomic data, which was measured for 134 (74 CD, 60 HF) of the 200 mice with MG data. In particular, we tested the expression of *Fasn* (fatty acid synthase), a key enzyme in hepatic lipogenesis, in relation to the F/B ratio, a combination which been shown as mechanistically related in the ceca of chickens [56]. In the CD group, *Fasn* expression showed a significant positive correlation with the F/B ratio (*r* = 0.52, *p* = 0.001), whereas no such association was observed in the HF group (*r* = 0.26, *p* = 0.16) (Figure 5 A). These findings suggest that the relationship between microbial composition and hepatic lipid metabolism may be modulated by dietary context, with stronger microbiota–host interactions under CD conditions. However, as previously discussed, phyla-level analyses of the microbiome may oversimplify their relationship to the host, and analyses at lower taxonomic ranks are necessary.

**Figure 5.**
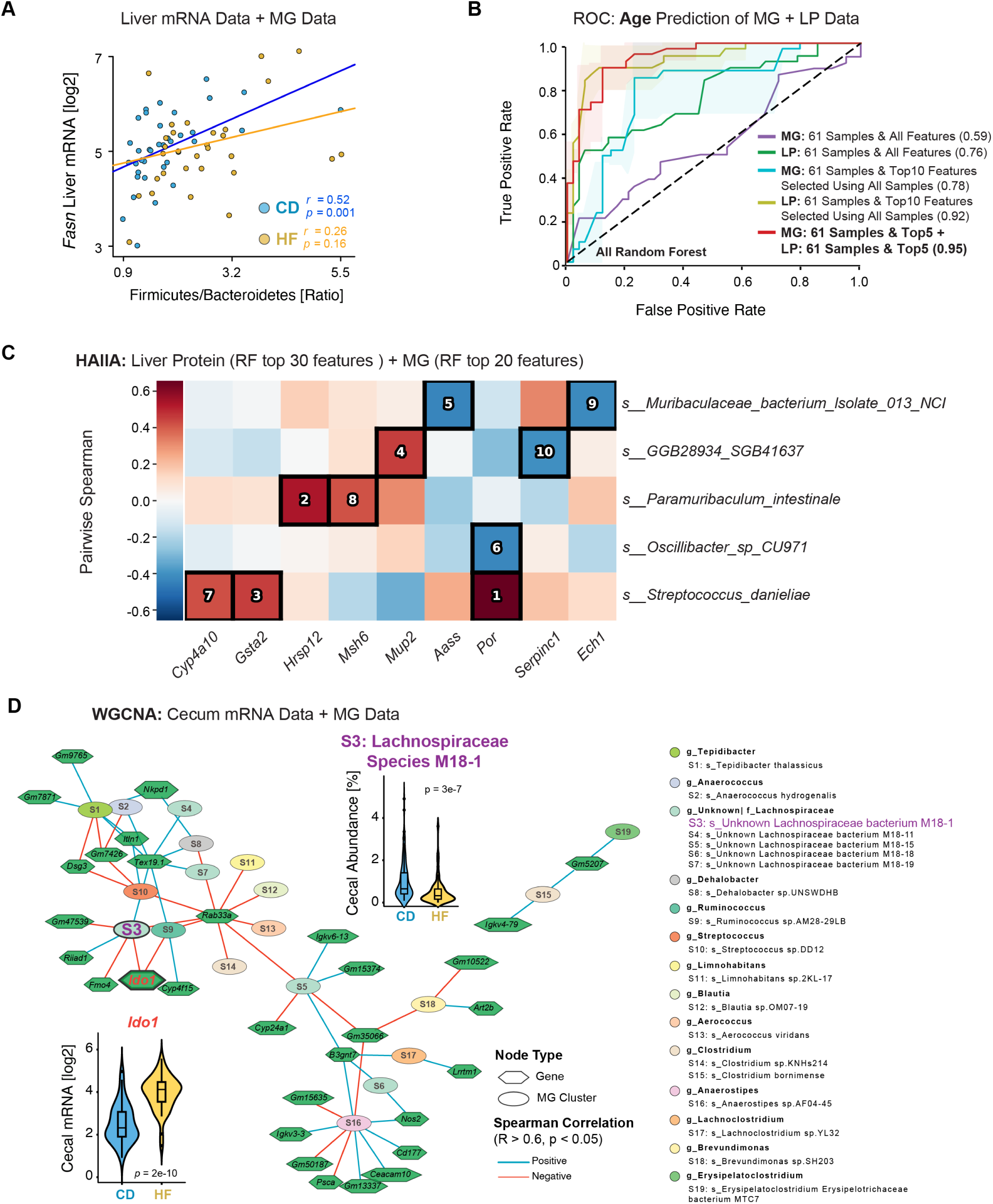
Integrated Multi-omics Analysis Reveals Age Prediction Performance and Microbial-host Interactions. **(A)** Correlation between Firmicutes / Bacteroidetes ratio and liver mRNA expression of *Fasn* stratified by diet group. (**B**) ROC curves for age prediction models based on MG and liver proteomic (LP) datasets, showing improved performance with integrated top features (AUCs up to 0.95). (**C**) Pairwise Spearman correlation heatmap between liver proteins and MG species. Black boxes mark significant correlations, with numbers showing their significance ranks. (**D**) Network visualization of host–microbiome correlations. Nodes represent either genes (hexagons) or species (ovals); edges indicate significant correlations (Rho > |0.6|, p < 0.05). Violin plots show the expression of *Ido1* and relative abundance of *Lachnospiraceae species M18-1*.

To further assess whether integrating multi-omics data could further enhance the performance of microbiome-based age classifiers, we focused on two datasets — MG species-level data and liver proteomics (LP) — which had the largest number of overlapping samples (n = 138) within our multi-omics collection. As a first step, we evaluated the predictive capacity of LP data alone. Using the same classification framework applied to the MG dataset, we trained classifiers on LP data from CD mice, binarized into Young (< 350 days) and Old (> 450 days) age groups. The RF model achieved an average AUC of 0.81 in 5-fold cross-validation (Figure S7 A, left panel) — equivalent to the results using MG alone (i.e. Figure 4 A). As with MG analysis, model AUCs improved marginally when moving from five to ten features (Figure S7 A, right panel), but no gain was observed beyond when using larger feature sets, despite that the liver proteome contains 3940 distinctly-measured proteins. Thus, like MG, a small subset of liver proteins was sufficient for robust age prediction. Consequently, we next selected the 138 overlapping samples between the MG and LP datasets and stratified them by diet. Within this set, 61 CD-fed mice fell into distinct Young (< 350 days, n = 25) or Old (> 450 days, n = 36) categories. These samples were then used to independently train age-prediction classifiers on each dataset (Figure 5 B). When using all available features from each dataset, model performance was modest (MG: AUC = 0.59, LP: AUC = 0.76), likely due to the small sample size relative to high feature dimensionality. To reduce the number of selectable features, several approaches can be undertaken, whether data-driven approaches (e.g. selecting high-variance features) or hypothesis-driven (e.g. selecting only MG features with annotated species names, or only selecting LP features within particular metabolic pathways). We therefore performed data-driven feature selection using the full MG and LP datasets, retaining only the top 10 features from each based on SHAP rankings from the top-performing models. Importantly, feature selection for MG and LP was performed using models trained on the full datasets (i.e. all 200 mice with MG data, and all 315 mice with LP data) rather than the reduced 61-sample subset, preventing information leakage or selection bias. By reducing the number of possible selectable features, we found a marked improvement in performance for both MG (AUC = 0.78) and LP (AUC = 0.92), demonstrating the utility of dimensional reduction in such analyses. Finally, we combined the top five MG features with the top five LP features to train an integrated classifier with both datasets together. This resulted in a compact 10-feature model, allowing direct comparison with the earlier models trained using the top 10 features from each individual dataset. This minimal multi-omics model achieved a striking AUC of 0.95, despite a relatively low sample size of 61 mice, suggesting that separate selection of key features across omics layers before multi-omic integration can substantially improve predictive performance. These results highlight the potential of compact, cross-modal biomarker panels to overcome limitations in multi-omics sample overlap, enabling more flexible study designs. The most predictive microbial and proteomic features identified here also offer promising targets for future mechanistic studies into host–microbiome interactions during aging.

Given the central role of the gut–liver axis in host metabolism, we next explored potential interactions between microbial and hepatic protein features identified through our machine learning models. Specifically, we used the HAllA framework [57] to assess associations between an expanded set (top 30 LP, top 20 MG) of features, selected by SHAP in the RF feature selection. The comparatively larger number of LP features was to reflect the greater feature dimensionality in the proteomic dataset (i.e. 3940 proteins), with the top 10 ranked associations displayed (Figure 5 C). We observed that *Streptococcus danieliae*, one of the most predictive MG features for age, exhibited strong positive correlations with three liver proteins — *Por*, *Gsta2*, and *Cyp4a10*. These cross-omics associations provide preliminary evidence for microbiome–host molecular crosstalk and highlight candidate gut–liver pathways for future mechanistic investigation.

Lastly, to further investigate cecum–microbiome interactions at the systems level, we constructed multi-omics co-expression networks (Figure S7 B-C, Figure 5 D) using Weighted Gene Correlation Network Analysis (WGCNA) based on cecal mRNA and MG species-level data. This analysis identified several modules containing tightly correlated host genes and microbial taxa. For example, cecal *Ido1* expression strongly associated with a group of HF-enriched species in the family Lachnospiraceae, including *Lachnospiraceae bacterium M18-1*. Previous studies have shown that *Ido1* knockout in mice alters gut microbial composition, and these mice exhibit significantly reduced white adipose tissue when fed a HF diet [58]. These findings support the intriguing possibility that, under HF conditions, *Ido1* may influence host metabolic outcomes through its interactions with specific species within the Lachnospiraceae family. Consequently, this integrated network approach links potential multi-layered signals between host and microbe, highlighting candidate functional relationships that warrant further mechanistic exploration. Together, these results underscore the value of combining statistical modeling, machine learning, and network-based strategies to extract meaningful biological insights from multi-omics data. They also establish a foundation for future hypothesis-driven studies aimed at dissecting causal microbiome–host interactions in the context of diet and aging.

## Discussion

Comprehensive studies analyzing the gut microbiota are showing new methodological approaches and diagnostic insights that researchers and clinicians can use to diagnose individuals’ current health concerns, and for prognosticating their future metabolic health. As with blood and urine, gut microbiome samples can be collected in the clinic longitudinally and minimally invasive. However, relatively limited annotation of bacterial species and their functions, combined with high amounts of variation in their abundances between individuals and over time, have slowed the general clinical application of metagenomic analysis. To understand key sources of variation across a highly-controlled population, in this study, we have focused on the ceca collected in 232 mice belonging to 43 strains of BXD mice over 1.5 years of age and in two different diets. Rapid improvements in sequencing quality and decreases in cost by orders of magnitude of cost have opened the door for large-scale sequencing studies on the composition and activity of the microbiome, moving beyond short-read 16S. It is now feasible for studies, like this one, to consistently identify and reliably quantify hundreds to thousands of taxa across hundreds of individuals.

Here, we generated and systematically integrated MG, MT, and host cecal mRNA data from a mouse population with controlled diversity to investigate how diet, aging, and genetic background interact and collectively shape gut microbial communities and host molecular responses. While genetic background and diet play a disproportionately large role in the microbiome composition and activity, age has a pervasive interactive effect — perhaps due to changes in immunity over time. At the taxonomic level, we quantified 614 microbial species in the MG dataset. Using supervised machine learning, we demonstrated that these profiles can predict key host phenotypes with high AUC-ROC performance: body weight (AUC = 0.92) and age (AUC = 0.84), underscoring the predictive capacity of the gut microbiome under diverse conditions, at least for highly-controlled mouse populations. Furthermore, through integrative modeling and prior feature selection between this dataset and our recently-collected liver transcriptome and proteome in the same mice, we were able to perform cross-tissue multi-omics analyses to obtain more reliable results with fewer selected features (e.g. 10 features to have AUC-ROCs of 0.95 for age). Given the dynamic responsiveness of the microbiome to environmental shifts and the convenience of sampling, this integrative approach offers powerful predictive capabilities for individualized assessments of future health trajectories, reflecting genetic and environmental contexts. Beyond improved predictive performance, this strategy effectively addresses the challenge posed by limited overlapping samples common in multi-omics studies, especially those with extended and heterogeneous sampling periods. By initially identifying robust biomarkers within individual omic datasets, researchers can subsequently leverage smaller, overlapping sample subsets for integrative modeling. This approach not only enhances study flexibility and resilience but is also highly applicable to human studies, where sample heterogeneity and logistical constraints often limit integrative analyses [59]. However, it must be emphasized that the massive role of genetic background and diet on MG and MT composition will result in challenges for applying this to non-inbred populations with little or no dietary control, like human cohorts.

Next, we sought to identify specific functional links to microbes and organismal function. Through an integrated analytical approach utilizing both MaAsLin3 and RF modeling, several microbial species consistently associated with aging emerged, notably *Streptococcus danieliae* and *Duncaniella dubosii*. *S. danieliae*, identified as strongly positively correlated with age in our analysis, aligns with prior findings from fecal 16S rRNA sequencing studies indicating that increased abundance of *Streptococcus* genera can accelerate aging [60]. These studies suggest that certain *Streptococcus* species could causally promote cellular aspects of aging, such as by elevating oxidative stress levels. Our findings raise the possibility that *S. danieliae* may represent a species-level candidate contributing to the observed age-associated microbial shifts. Conversely, *D. dubosii* showed a significant negative correlation with aging, highlighting its potential protective role. Prior studies have linked microbial modulation of tryptophan metabolism to longevity and age-related disease resistance [61], and intriguingly, the survival benefits conferred by tryptophan were found to be microbiota-dependent. While its specific role in aging remains to be determined, our findings nominate *D. dubosii* as a candidate taxon for further investigation into microbiome-based mechanisms supporting healthy aging. Furthermore, several of our analyses identified unannotated, merged cluster of genera within the Lachnospiraceae family as associated with dietary effects and functional divergences in bacterial activity (e.g. Fig Figure 2 D, Fig Figure 3 B). To attempt to disentangle this cluster of families, we processed the MG data using the Kraken2 pipeline [62] to look at all genera within the Lachnospiraceae family through a separate read mapping tool (data in Table S9).

Our network analysis using WGCNA identified significant associations between cecal expression of *Ido1* and specific Lachnospiraceae, particularly *Lachnospiraceae bacterium M18-1*. Notably, under HF conditions, we observed a significant increase in *Ido1* expression alongside a marked reduction in the abundance of several Lachnospiraceae species. This inverse pattern was reflected in a negative correlation within the co-expression network. Although no prior studies have directly examined interactions between *Ido1* and individual species within the Lachnospiraceae family, existing literature provides a plausible mechanistic link. Members of Lachnospiraceae are core constituents of the gut microbiota and are major producers of SCFAs such as butyrate [63]. SCFAs have been shown to inhibit the expression of *Stat1*, thereby downregulating IFNγ-induced transcription of *Ido1* [64]. Our findings are consistent with this model, suggesting that a HF-induced decline in SCFA-producing microbes may relieve inhibitory control over *Ido1*, leading to its upregulation. Furthermore, *Ido1* knockout has been associated with altered microbiota composition under HF conditions [58], pointing to a bidirectional feedback between host tryptophan metabolism and microbial ecology. These observations underscore a potential regulatory axis between microbial SCFA production and host immune-metabolic responses via the *Ido1* pathway.

Several limitations of our study should be acknowledged. Firstly, the sample size per strain restricted detailed analysis of the genetic impact across diverse BXD strains. While the overall effect of genetic background could be clearly determined due to the large number of strains (43), only 7 strains had 10 or more individuals analyzed. Thus, the study is not well-powered to determine effects on the microbiome that are specific to individual outlier strains. Secondly, many identified microbial taxa of key functional interest, such as *“GGB28815 SGB41470”*, remain poorly characterized and their cultivation conditions are unknown, complicating downstream functional validations through traditional microbiological methods. Further development in culturomics and advanced molecular techniques are required to overcome these barriers. Additionally, our study relied on cross-sectional sampling at discrete age points, precluding longitudinal analyses that could offer deeper insights into microbiome dynamics and causal inference. Although inbred strains have been used, variation between biological replicates of up to 30% of overall variation creates challenges for establishing definitive timelines of change, even for microbes with highly-significant associations with age. Lastly, numerous other independent variables that were not in the scope of this study can have a major effect on the microbiome. For instance, sex has a pronounced effect on the gut microbiome [65] and on metabolic traits. However, 97% of the samples in this study are from female mice due to logistical challenges with aging male mice (particularly fighting).

In conclusion, our integrative multi-omics approach highlights the influence of dietary patterns and aging on the gut microbiome composition and functional landscape, identifying key microbial taxa and metabolic pathways intricately linked to the host organism’s health. Our results further underscore the remarkable utility of microbiome profiling combined with machine learning for reliable host age prediction, underscoring its potential application as a reliable biomarker for biological aging and personalized health monitoring. Furthermore, still nearly 70% of specifically-identified species in the shotgun metagenome sequence data have placeholder names (e.g. *SGB62427*) as they await more specific functional analysis and formal naming. Multi-omics integration and the study of how these species vary according to key experimental variables guides future studies into these bacteria and their mechanistic relationships with metabolism (e.g. that *SGB40300* tracks consistently with age, across two different dietary cohorts). Together, these findings provide a robust foundation for future mechanistic explorations and translational applications aiming to harness microbiome insights for metabolic health interventions and age-related health assessments.

## Methods

### Mice Sample Collection and Processing

All animal handling and tissue collection were done in the scope of our prior study [17], but it is briefly recapitulated here. 662 animals from 58 strains of the BXD family were sacrificed at specific ages (200, 365, 540, or 730 days, each with a standard deviation of about 30 days), of which cecal tissue was retained for 597 individuals from 56 strains (Table S1). The aging colony was 95% females due to methodological challenges of aging large groups of males (e.g. fighting during regroupment). Diets were either Harlan Teklad 2018 (CD; 24% calories from protein, 18% from fat, 58% from carbohydrates) or Harlan Teklad 06414 (HF; 18.3% calories from protein, 60.3% from fat, 21.4% from carbohydrates). Animals were removed from the aging colony the night before sacrifice but retained access to food and water. Animals were anesthetized between 9am and noon with tribromoethanol and perfused with PBS. Cecal tissue containing fecal matter was harvested within about 7 minutes of perfusion, then immediately frozen in liquid nitrogen and stored in -80°C freezers. 274 samples were selected from the full set of 597 ceca to measure strains that had tissues collected in both diets and at multiple ages. The female mice represented both dietary conditions across the 44 strains, whereas male samples were selected only for B6 mice fed the HF diet, due to the sex imbalance in the initial aging study. The subset of 274 samples was selected with the goal of balanced sample size of diets and age groups - except for the fourth age group, which was reduced as several strains do not typically live to 2 years of age. After accounting for balanced diets and ages, half of the samples were selected to give high coverage of specific strains at multiple diets and timepoints (e.g. 19 C57BL/6 mice were selected) and the remaining half of samples was selected to increase the breadth of genotypes analyzed to cover the full set of 56 strains as closely as possible.

For these 274 samples, the mixed cecum and cecal contents tissue samples were transferred from a -80°C freezer to dry ice. Prior to grinding, the mortar and pestle were pre-cooled with liquid nitrogen. To ensure consistent extraction across all samples, the entire frozen cecal tissue was homogenized in liquid nitrogen using the pre-cooled mortar and pestle until fine powders were obtained. After the sublimation of the liquid nitrogen, the powder was transferred to new tubes using a round spatula scraper, all while remaining at around -196°C. A ^_^30 mg aliquot of the mixed cecum and cecum content powder was taken both for DNA and RNA extraction, with all 3 aliquots (i.e. for DNA, for RNA, and remainder) stored in a -80°C freezer until extraction. MG, MT, and cecal mRNA samples were extracted and sequenced in two temporally distinct batches - November 2022 and November 2023 - by the same person using the same protocol. After sample preprocessing, 232 of the initial 274 samples remained across 43 out of 44 strains, of which 225 were from female mice and the remaining 7 from males, similar to the sex distribution in the original aging colony (Table S1, second sheet). The loss of 42 samples was for three reasons: improper freezing of the tissue, incorrect tissue in the tube, or failure to extract good-quality DNA or RNA from the sample. An overview of all datasets and their sample distributions across diet, age, and strain is provided in Table 1. Then, 10 of the 232 mice were measured multiple times across the two batches (suffix “_r” or “_n” in Table S1, second sheet), for either re-sequencing of the exact same DNA sample (r) or a full re-extraction, purification, and sequencing of DNA from a powdered tissue aliquot from the same initial sample (n). These samples were used as a sanity check for possible batch effects caused by sample preparation or sequencing, as shown in the right panels of Figure 2 D and Figure 3 A.

To note, while during the initial sample collection, cecal tissue can be opened up and cecal contents separated out and frozen separately. For this study, which focused on MG or MT analysis, collecting the frozen cecum as a whole can provide some analytical benefits (no batch effects caused by procedural differences in scraping out the contents of a cecum) and practical benefits (no special training is necessary if samples are collected by collaborators, and no significant extra time is required during the sacrifice to collect a whole cecum). However, extraction, separation, and measurements of the (meta-)proteome or (meta-)metabolome would not be feasible with the above protocol.

### Optimization of DNA Extraction Protocol

For DNA extraction, a modified extraction method employing the Wizard Genomic DNA Purification Kit (catalog number: A1120) was used to simultaneously extract DNA from both the host and the gut bacteria in mixed samples of cecum and feces. ^_^30 mg of tissue powder was suspended in 480 µL of 50 mM EDTA and homogenized with stainless steel milling beads using an Oscillating Mill MM400 (Retsch) at 25 Hz for 1 min, then incubated at 37°C for 60 min. After the homogenates were centrifuged for 2 min at 500 g, the supernatant was transferred into a fresh tube. This supernatant was then centrifuged for 2 min at 16,000 g. After centrifugation, the supernatant was removed and the pellets were collected. The pellets were lysed by adding 600 µL of chilled Nuclei Lysis Solution, gently mixed by pipetting, incubated at 80°C for 5 min, and subsequently cooled to room temperature. Next, 3 µL of RNase Solution was added, and the mixture was incubated at 37°C for 45 min before cooling to room temperature. Protein precipitation involved adding 200 µL of Protein Precipitation Solution (from the same Wizard kit), followed by vortexing and a 5 min incubation on ice. The sample was then centrifuged at 13,000 × g for 3 min. Subsequently, the supernatant was transferred to a fresh tube with 600 µL of isopropanol for DNA precipitation and mixed gently by inversion. This was followed by a centrifugation at 16,000 × g for 2 min to remove the supernatant. Next, 600 µL of 70% ethanol was added, mixed, and the sample was centrifuged again for 2 min at 16,000 g. Ethanol was removed by aspiration, and the pellet was air-dried completely, which took roughly 15 min. The DNA pellet was resuspended in 100 µL of Rehydration Solution for 1 hour at 65°C.

As we are performing shotgun sequencing on the MG samples (rather than 16S), this step was necessary to remove most of the mouse DNA content of each sample, which was not of interest given that the BXDs are inbred and have recently been fully sequenced [66]. Thus, to remove mouse DNA for MG sequencing, we utilized the Wizard DNA Clean-up System for size-exclusion following the manufacturer’s protocol. To check the final ratio of mouse-to-microbe DNA in the purified sample, the concentration and purity of each DNA sample were measured using a spectrophotometer (Nanodrop 2000C, Thermo Scientific). We established a method for evaluating the proportion of gut bacterial DNA using RT-qPCR with the ratio of *Gapdh* primers (Forward: CAAGGAGACCTCAAGGTCAG; reverse: GTATGCACCTCACAACGCCATG) for the eukaryotic content compared to the universal prokaryotic 16S rRNA primers (Eurogentec, Forward: AAACTCAAAKGAATTGACGG; reverse: CTCACRRCACGAGCTGAC). Host DNA and intestinal bacterial DNA were extracted separately from liver and true fecal samples (i.e. using excreted feces, rather than cecal content) and then diluted to a concentration of 10 ng/µL. Subsequently, different ratios of host DNA and gut microbiota DNA were mixed to serve as controls. The concentration of all samples was adjusted to 10 ng/µL for use in RT-qPCR assays. RT-qPCR was performed using the LightCycler 480 Instrument II (Roche). Each reaction had 2.5 µL of cDNA, 2.5 µL of primer mixture (forward and backward primers) (2 µM) and 5 µL of the iQ SYBR Green Supermix (BIO-RAD). The PCR reaction conditions were as follows: an initial denaturation step at 95°C for 10 min, followed by 40 cycles of denaturation at 95°C for 15 seconds and annealing/extension at 60°C for 1 min. The reaction was subsequently cooled to 20°C for 1 hour. Based on the Crossing Point (Cp) values obtained from the RT-qPCR results, we defined a %Expression index to estimate gut microbiota-to-host DNA ratios and used it to filter samples with higher bacterial DNA content. The %Expression index was defined as

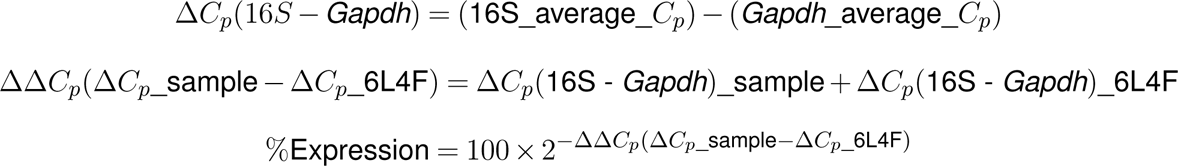

Where 16S_average_Cp is the average Cp value from three technical replicates of each DNA sample extracted from mouse cecal contents, using universal prokaryotic 16S rRNA primers; and *Gapdh*_average_Cp is the average Cp value from the same samples using primers targeting the mouse reference gene *Gapdh*. The value of `Δ`C*_p_*_6L4F refers to the Cp difference calculated from a reference DNA mixture composed of 40% fecal DNA and 60% liver DNA (mixed at equal concentrations), which serves as a normalization control across samples. Samples with a gut microbiota DNA proportion estimated to be greater than 40%, based on the Cp values of the controls, were retained for MG sequencing (Figure S1 B-C). This cutoff was made for financial considerations, considering only that if a sample is e.g. 70% mouse DNA, then a large majority of reads (i.e. those derived from the mouse genome) would be discarded and result in lower effective read depth for such samples.

### RNA Extraction Protocol

For both MT and cecal mRNA extraction, the same ^_^30 mg of tissue powder was suspended in 1 mL ice-chilled TRIzol (Thermo Scientific) and homogenized with stainless steel milling beads in an Oscillating Mill MM400 (Retsch) at 25 Hz for 30 seconds, followed by a phase separation using 200 µL of chloroform, and the mixture was vortexed before being left at room temperature for another 2 min. The mixture was centrifuged at 12,000 g for 15 min at 4°C, the upper aqueous phase was then transferred to a new RNase-free tube. For RNA precipitation, 400 µL of isopropanol was added and thoroughly mixed by pipetting, followed by incubated at room temperature for 2-5 min. This was followed by centrifugation at 12,000 g for 10 min at 4°C, with the supernatant was discarded. The RNA pellet was washed with 1 mL of 75% ethanol, centrifuged at 7,500 g for 5 min at 4°C, and the supernatant was subsequently discarded. Finally, the RNA pellet was air-dried at room temperature for 10 min before being resuspended in 50 µL of RNase-free water. For RNA purification, the RNeasy MinElute Cleanup Kit (catalog number: 74204) was utilized following the manufacturer’s protocol (Qiagen, March 2016 version). For RNA quality assessment, the concentration and purity of each RNA sample were measured using a spectrophotometer (Nanodrop 2000C, Thermo Scientific). RNA quality was evaluated using an Agilent RNA 6000 Nano Kit and Agilent 2100 Bioanalyzer. Given that the ‘ideal’ RNA samples contain both host RNA (for cecal mRNA) and gut bacterial RNA (for MT), the RNA Integrity Number (RIN) is not usable for these mixed samples. Only samples exhibiting four visually-distinct peaks were selected for subsequent sequencing (Figure S1 D).

### Sequencing and Profiling

#### Sequencing

The mRNA sequencing libraries were prepared using the TruSeq Stranded mRNA Library Prep Kit (Illumina, catalog number: 20020594). Sequencing was performed on the Illumina NextSeq 500 system, utilizing single-end sequencing with a read length of 75 base pairs. MT sequencing was conducted after library preparation using the Zymo-Seq RiboFree Total RNA Kit (catalog number: R3003), and sequencing was carried out on the Illumina NextSeq 2000 system with 2×150 bp paired-end reads. MG sequencing used the Illumina HiSeq 1000, with roughly 25 million reads per sample, using 2×150 bp paired-end reads. Quality assessment of the MG and MT sequenced reads, along with their corresponding bases, was conducted using FastQC v0.12.1 (RRID:SCR 014583). In our case, all reads and bases were within an acceptable quality range, with Phred scores consistently exceeding 30, indicating high-quality sequencing data.

#### Cecal mRNA Data Processing and Analysis

For gene expression analyses, the cecal mRNA sequencing data were aligned and quantified to generate per-gene unique read counts using the R package `Rsubread` v2.12.3 (RRID:SCR 016945) [19], resulting in a raw gene count matrix (Table S2). This raw count matrix was used for DEG analysis in `DESeq2` v1.44.0 (RRID:SCR 000154) [67] to identify transcriptional changes associated with diet and age, and the resulting log_2_(fold changes) were used for GSEA. In parallel, raw counts were also converted into a TPM (Transcripts Per Million) matrix using the package `tximport` v1.26.1 (RRID:SCR 016752) [68], followed by log transformation as `log`_2_(TPM+ `1)` (Table S3) for use in machine learning models and WGCNA. This dual processing strategy enabled robust DEG identification while facilitating pathway-level interpretation and comparison across samples. For the volcano plots in Figure 1, the -10log(raw p-values) were used for graphic visualization, but note that the adjusted p-value cutoff of 0.05 was used for statistical interpretation of the data. Genes with raw p-values above 0.05 were not plotted in the volcano plots (i.e. genes with y-axis values less than 1.3), as vector graphic images slow down substantially with tens of thousands of objects. ORA and GSEA were performed using the `clusterProfiler` v4.2.2 (RRID:SCR 016884) [69] package in R, based on the KEGG mouse pathway database (*organism* = “mmu”). In ORA, the “rich factor” is a normalized metric for the ratio of how many genes in that pathway are differentially expressed compared to the overall gene set size of that pathway, e.g. a rich factor of 0.25 means 25% of genes in that pathway are differentially expressed. The functions `enrichKEGG` and `gseKEGG` were used to identify significantly enriched pathways among differentially expressed genes. Gene annotation was supported by `org.Mm.eg.db` (v3.20.0).

#### MG and MT Data Processing

All raw paired-end MG and MT sequence reads were quality-processed using KneadData v0.10.0 [36]. Sequence trimming was performed using KneadData, which ran Trimmomatic v0.39 (RRID:SCR 011848) to trim regions where base quality fell below Q20 within a 4-nucleotide sliding window, and to remove reads that were truncated by more than 50% (parameters: SLIDINGWINDOW:4:20, MINLEN:50). Additionally, KneadData was used to run Tandem Repeat Finder (TRF) v4.09.1, to locate and remove reads characterized by tandem repeats. Bowtie2 v2.4.2, also employed within KneadData, was used to deplete host-derived sequences from the MG data using the mouse (C57BL/6, RRID:IMSR JAX:000664) reference database, and from the MT data using both the mouse reference database and the SILVA Ribosomal RNA reference database (RRID:SCR 006423). After processing with KneadData, the remaining non-host reads were used as input to obtain the taxonomic profiles of the samples using MetaPhlAn v4.0.6 (RRID:SCR 004915) and the CHOCOPhlAn (mpa_vOct22_CHOCOPhlAnSGB_202212) database. Functional profiling was subsequently obtained through the HUMAnN v3.8 (RRID:SCR 014620) workflow. To note, different MG and MT alignment pipelines can still generate quite divergent estimates of microbiome abundances, particularly at lower taxonomic levels where read mapping is more challenging and reference databases are still changing more quickly. We highly recommend particularly interested readers to download the raw data (on NCBI’s SRA, reference PRJNA953833) and process the sequence data on their own pipeline. For those interested in a cursory look at the differences between read mapping on different pipelines, we have included the MG data mapped using the Kraken2 pipeline (RRID:SCR 026838) [62] in Table S9.

### Miscellaneous Bioinformatics and Statistical Analysis

All analyses were performed using R 4.2.2. For exploratory data visualization, differential expression analysis, statistical modeling, and multivariate analyses (e.g., ANOVA and PCA), we used standard R packages including `ggplot2` (RRID:SCR 014601) [70] and `vegan` [71]. Comparisons of abundance between two dietary groups was performed by Welch’s t-test. ANOVA was used for comparison of multiple groups simultaneously, with Tukey post-hoc tests used afterwards to test for differences between groups (e.g. Figure 1 D). In figures with linear correlations between two variables, we used either Pearson correlation or Spearman correlation coefficients. Pearson correlation (represented by r) was used when all points could be clearly visualized in the plot and no outliers were detected. Pearson correlation (represented by R) was used for generating networks (e.g. Figure 5) and for testing the covariance of variables with high variation or outliers (e.g. correlating MG to MT abundances). Data normality was checked with Shapiro-Wilk test in R (shapiro.test), with W values above 0.80 considered normal, regardless of the p-value. False discovery rate (FDR) was controlled using the Benjamini–Hochberg (BH) procedure either manually with the p.adjust() function in R, or built-in as part of package analyses such as with MaAsLin3. Features with FDR < 0.05 (adj.p < 0.05) were considered statistically significant. To identify modules of co-varying host genes and microbial taxa, we performed WGCNA using the R package: `WGCNA` [72]. For the network in Figure 5, cross-omics edges were defined by Spearman correlation coefficients |*R*| > 0.6 and FDR-adjusted *p* < 0.05. Significant gene–microbe interactions were exported as edge lists and visualized using Cytoscape (v3.10.3, RRID:SCR 003032) [73]. In Figure S1, we used Spearman correlation to correlate the relative abundances of the same 38 highest-detected taxa in the MG and MT data across the 48 mice with measurements on both omics layers. For the negative control for the cross-omics correlation of MG versus MT abundances, we first subset to take the 38 most-abundant genera at the MG and MT level. These 38 genera at the MG level were then correlated to a randomly-selected one of the same 38 genera at the MT level, without repetition and ensuring that the randomized index did not match the original index. Regarding multi-omics overlap, 48 exact mice had overlapping data at the MG-MT-cecal mRNA dataset. This is due to the order of operations in our study design: the MT and mRNA data were generated first due to their technical challenge, and this used the entirety of our cecal tissue for around 25 samples. For 19 of these samples, we had biological replicates (i.e. the same strain, age, sex, and diet), so those biological replicates were selected for MG analysis. Thus in total 67 *cohorts* have MG-MT-mRNA data, but we have used only the 48 perfectly-aligned *mice* for simultaneous analysis of multi-omics layers.

### Multivariable and Cross-Omics Association Analysis

To identify microbial features associated with metadata variables such as diet, age, and strain, we applied MaAsLin3 (v1.10.0) [46], a multivariable linear modeling framework suitable for compositional microbiome data. Relative abundance tables (species- or genus- level) from MetaPhlAn4 [37] were were first normalized using total sum scaling (TSS) and then log-transformed. Features present at a relative abundance of at least `1` × `10`*^−^*^4^ in ≥ `10%` of samples were retained. Diet- and age-associated analyses were performed using separate MaAsLin3 models, with the target variable set as the fixed effect and the other variables set as random intercepts. For instance, for diet associations, we used the model Abundance ∼ Diet + Age + Strain. For age, we used Abundance ∼ Age + Diet + Strain. In MaAsLin3 output, “abundance” refers to the normalized relative abundance of a microbial feature in those samples where it is detected, while “prevalence” denotes the proportion of samples in which the feature is present (i.e. non-zero). The estimated coefficients summarize how each metadata variable relates to these measures after adjusting for other covariates.

To explore cross-modal associations between microbial species and host liver proteomic features, we used the HAllA framework [57]. HAllA identifies statistically significant associations between high-dimensional datasets by combining hierarchical clustering and association testing across feature blocks. For downstream visualization, we ranked the resulting association blocks by HAllA’s hierarchical association score and plotted the top ten blocks using the `hallagram` tool. In these hallagrams, each colored cell represents the strength and direction of the pairwise Spearman correlation between one microbial species and one liver protein within the corresponding cross-omics block.

### Supervised Learning and Model Evaluation

All prediction models were implemented using Python 3.9.18 (RRID:SCR 024202) with standard open-source libraries, including `pandas` v2.1.4 (RRID:SCR 018214) [74], `numpy` v1.23.5 (RRID:SCR 008633) [75], `scikit-learn` v1.3.0 (RRID:SCR 002577) [76], `SHAP` v0.48.0 (RRID:SCR 021362) [77], and `matplotlib` v3.8.0 (RRID:SCR 008624) [78]. We trained multiple supervised classifiers including RF, XGB, Logistic, and DNN to predict host phenotypes (e.g., age and body weight) based on gut microbial composition. Model performance was evaluated using five-fold cross-validation, and averages across folds were used to generate ROC curves. Feature importance scores were extracted from tree-based models (RF and XGB) to rank the contribution of individual microbial species or protein features. Feature ranking was performed using models trained on the full MG (n = 200) and LP (n = 316) datasets, ensuring that feature selection was independent of the reduced sample subsets used in downstream multi-omics integration. To complement these model-derived rankings and improve interpretability, we computed SHAP (SHapley Additive exPlanations) values to quantify each feature’s contribution to individual predictions. The top 20 features were visualized using SHAP beeswarm plots, in which the horizontal position reflects the magnitude and direction of the SHAP value while the color gradient represents feature abundance across samples. To mitigate overfitting and improve interpretability, prior feature selection was applied based on model-derived importance scores. To further capture potential nonlinear or combinatorial effects between microbial features, polynomial feature interaction terms were generated using the `PolynomialFeatures` module from `scikit-learn`. In multi-omics integration analyses, classifiers were trained using top-ranked features from each omics layer and evaluated using a compact, 10-feature integrated model. Model optimization was guided by cross-validation accuracy and AUC metrics across independent data batches.

## Data Availability

The raw sequence data are available in the NCBI Bioproject repository under the accession number PRJNA953833 (data will be released upon article publication). All processed data tables for the MG, MT, and cecal mRNA are available in nine supplemental tables (S1-S9). All tables are also available at the GitHub repository together with the project code (https://gitlab.com/uniluxembourg/lcsb/gene-expression-and-metabolism/2025_agingcecum_mgmtmrna_zzhou).

## Acknowledgements

We would like to thank Kurt Lamour for assistance with DNA sequencing. We are grateful to Xiaowei Zhan, Susheel Bhanu, and Julien Schleich for their valuable advice on data processing and analysis. We also thank Jack Haverty for his help during the early stages of the project. Rob Williams and Suheeta Roy from the University of Tennessee Health Science Center are acknowledged for hosting the aging mouse colony and their support in the organization of cecal sample collection. The mRNA and MT sequencing experiments presented in this paper were carried out at the LCSB sequencing platform (RRID: SCR_021931) at the University of Luxembourg. The processing of sequencing raw data presented in this paper were carried out using the HPC facilities of the University of Luxembourg [79].

## Author Contributions Statement

E.W. and P.W. were responsible for the initial conceptualization and study design. E.W. supervised and provided guidance throughout the research process. Z.Z. and E.W. wrote the main manuscript text. Z.Z. prepared all figures and tables, with individual panel input and figure editing from E.W. Z.Z. optimized protocols for simultaneous extraction of host cecal and gut microbial DNA and RNA for sequencing. Z.Z. prepared all samples used in this manuscript. R.H. performed RNA sequencing. A.A., E.P., B.B., and T.K. assisted with various practical aspects of the experiments and exploratory analyses. S.N. assisted with analysis and writing for the revision. All authors reviewed the manuscript.

## Ethics Declarations

### Ethics approval and consent to participate

The mice from this study were approved by the University of Tennessee Health Science Center’s committee for animal care and use, following AAALAC guidelines and with regular inspections by independent veterinarian oversight.

### Consent for publication

Not applicable.

### Code Availability

Codes and scripts developed in this study are all available at the GitHub repository (https://gitlab.com/uniluxembourg/lcsb/gene-expression-and-metabolism/2025_agingcecum_mgmtmrna_zzhou).

### Supplemental Information

Supplemental Information contains seven figures and nine supplemental tables with the metadata for the cecal samples, and the sequence result data for the MG, MT, and cecal mRNA.

## Supplementary Information

**Figure S1.**
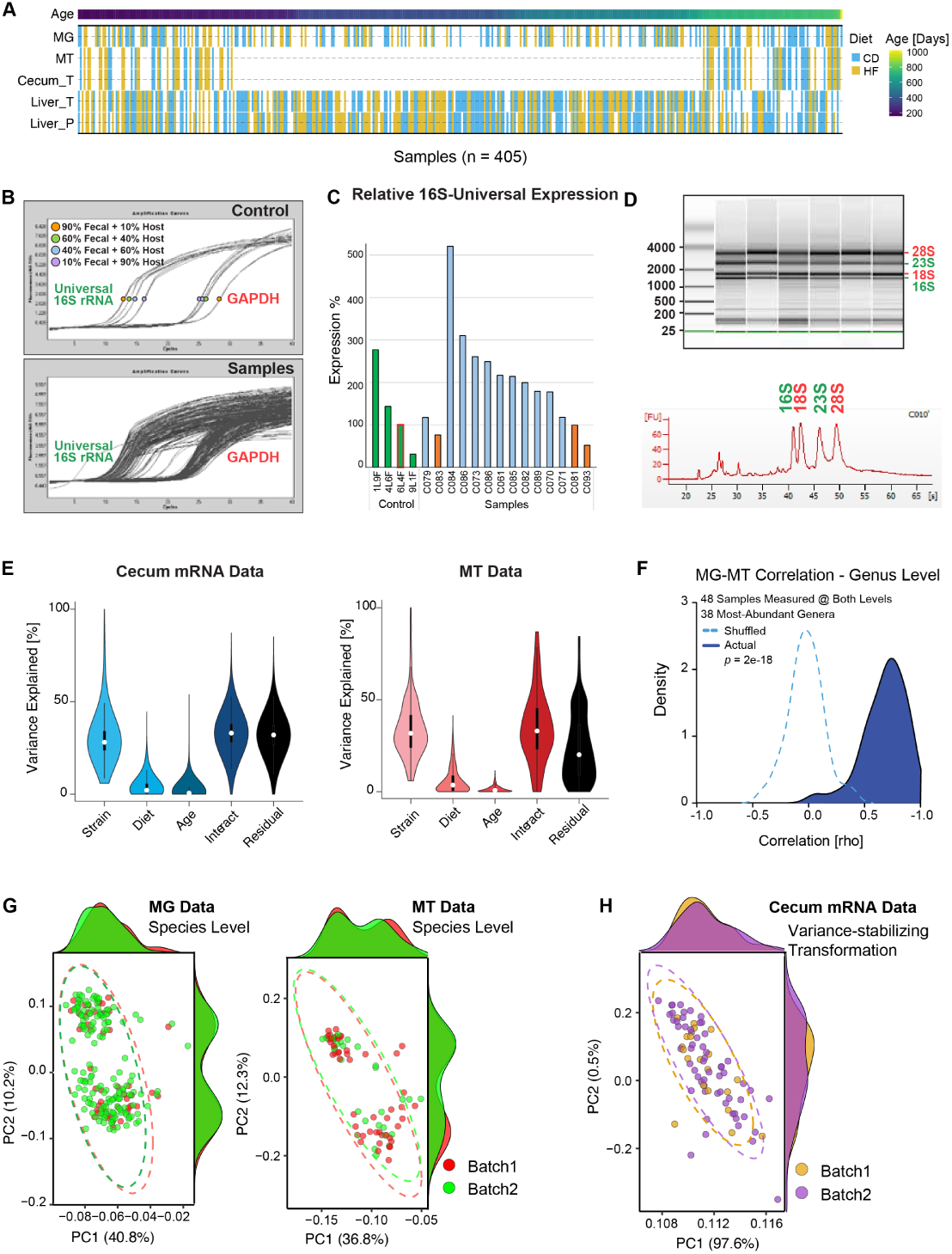
Establishment of Experimental Methods and Quality Control. (**A**) Overview of sample availability across all omics layers. The heatmap displays the presence or absence of each data type for all 405 samples, including metagenomics (MG), metatranscriptomics (MT), cecal transcriptomics (Cecum_T), liver transcriptomics (Liver_T), and liver proteomics (Liver_P). Top annotations indicate age (days). For specific sample overlaps in the cecal data, refer to the supplementary tables. (**B**) In order to check whether purified MG samples had high microbiome content and low mouse content, RT-qPCR primers were used to check for the relative abundance in both sources of DNA. (**C**) Samples with gut microbial DNA content estimated to be above ∼40% were selected based on gradient controls; a control sample was generated by mixing DNA extracted from the liver with DNA extracted from fecal samples in different ratios. (**D**) For RNA, Bioanalyzer results were used to select samples with four clear RNA bands, indicating abundance of both mouse and microbiotal RNA, which were then separated. (**E**) ANOVA analysis of cecal mRNA and MT data showing variation explained according to the study variables. (**F**) Correlation analysis indicates a strong correlation between the MG and MT data for the most abundant 38 genera; “shuffled” indicates 38 genera at the MG level correlated with 38 randomly-selected genera at the MT level as a negative comparison. (**G**) PCA plots of MG and MT at the species level show no dominant batch effect according to the two extraction batches (performed in Nov 2022 and Nov 2023 for MG data, and Dec 2020 and Nov 2022 for MT data). The two distinct clouds within MG and MT data show the dietary groups. (**H**) PCA based on variance-stabilized cecal mRNA expression data shows that cecal samples also do not cluster by batch, nor is the effect of diet immediately apparent.

**Figure S2.**
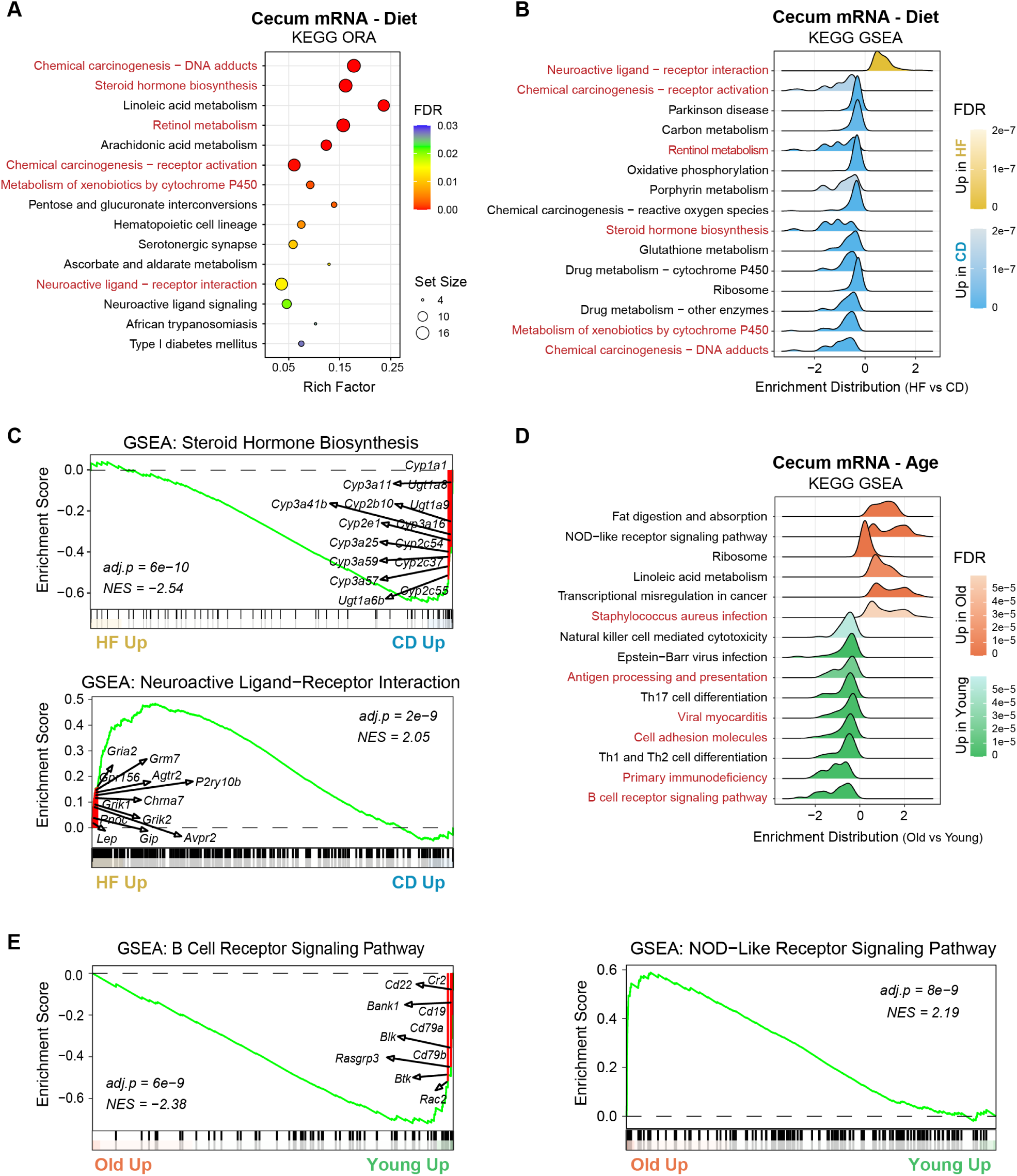
Differential Pathway Enrichment in Cecal Transcriptomes by Diet and Age. **(A)** KEGG ORA results showing the top 15pathways affected by diet in cecal mRNA. Bubble size indicates the number of DEGs per pathway; bubble color represents FDR; the x-axis shows Rich Factor. **(B)** KEGG GSEA for the same diet comparison, displaying the top 15 pathways ranked by FDR. **(C)** Two select GSEA enrichment plots for diet: steroid hormone biosynthesis (upper; enriched in CD) and neuroactive ligand–receptor interaction (lower; enriched in HF). Marked genes are DEGs present in the respective pathways. **(D)** KEGG GSEA results showing the top 15age-associated pathways. (E) Two select GSEA plots for age: B cell receptor signaling (left; enriched in Young) and NOD-like receptor signaling (right; enriched in Old).

**Figure S3.**
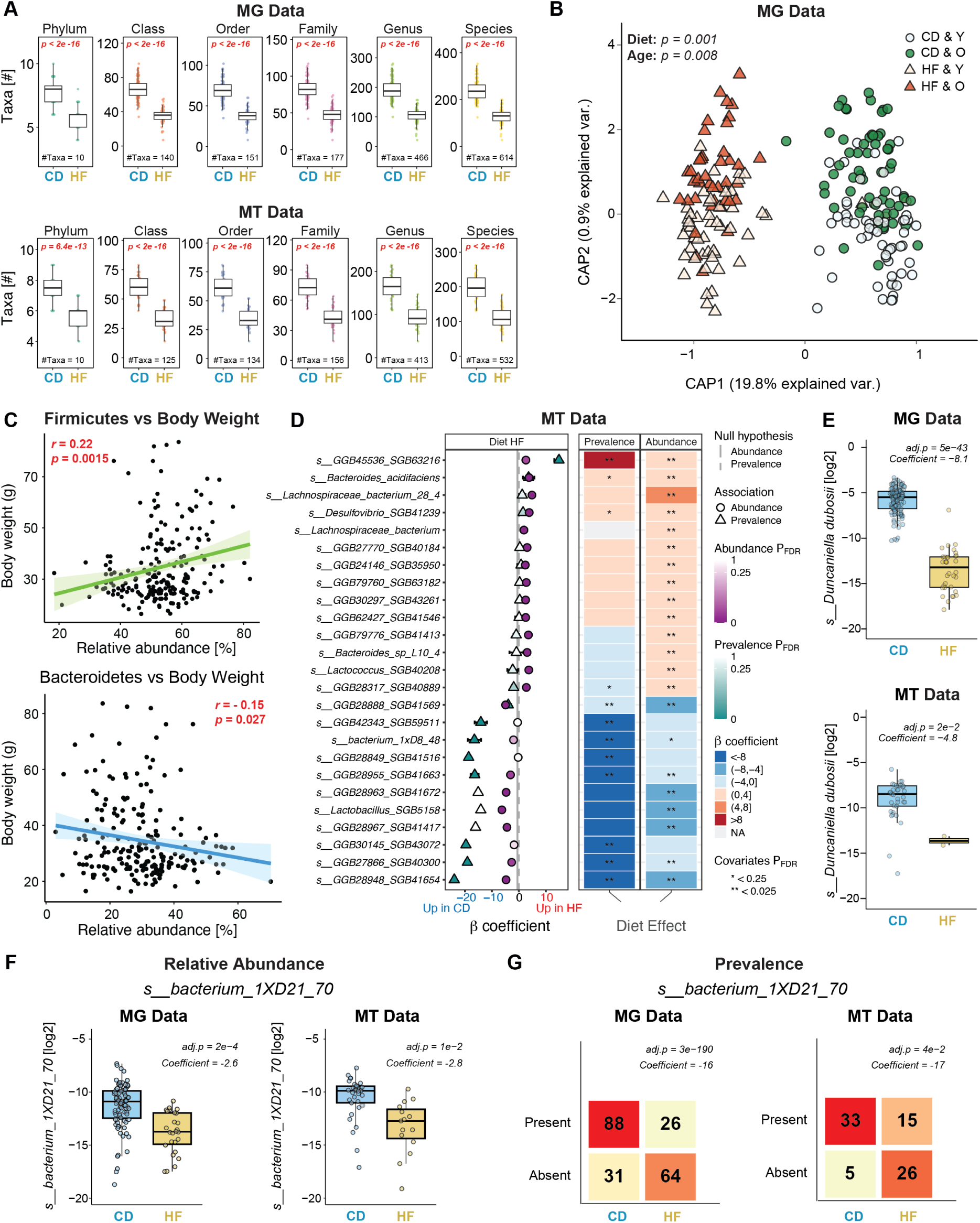
Supplementary Analysis of MG and MT Data. **(A)** Total number of detected gut microbial taxa at all taxonomic levels (binary detection cutoff, with even 1 read considered a detection), separated by diet. HF significantly reduced microbial richness at each taxonomic level. **(B)** Distance-based redundancy analysis(dbRDA) based on Bray–Curtis dissimilarity showing gut microbial community structure across diet and age (CD-Young, CD-Old, HF-Young, HF-Old). *P*-values from PERMANOVA (999 permutations). **(C)** Firmicutes and Bacteroidetes phyla are significantly associated with body weight, across all mice. **(D)** MaAsLin3 plot and heatmap of MT data, showing the species affected most by diet. **(E)** The relative dietary abundance of *Duncaniella dubosii* **(F)** and of *Bacterium 1XD21 70* across different diets in both MG and MT data. **(G)** The binary prevalence metric for the same two species in the two datasets.

**Figure S4.**
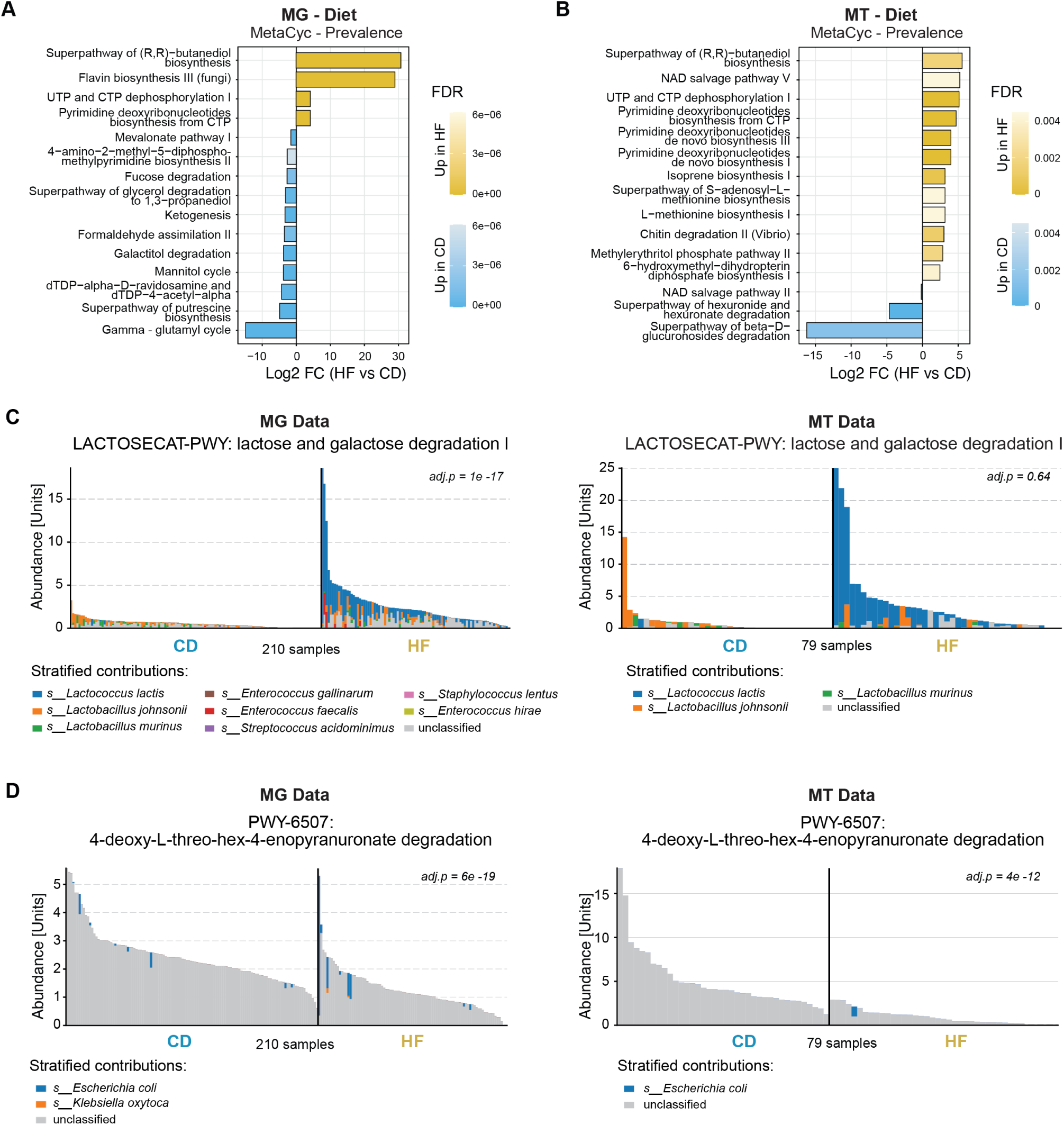
Supplementary Details of Diet-associated Changes in Microbial Functional Profiles. **(A)** Top 15 MetaCyc pathways showing significant differences according to diet in pathway prevalence using MG data and **(B)** MT data **(B)**. **(C-D)** Relative abundance distributions of the lactose and galactose degradation I pathway (LACTOSECAT-PWY) **(C)** and the 4-deoxy-L-threo-hex-4-enopyranuronate degradation pathway (PWY-6507) (D) in the MG and MT datasets, stratified by major contributing species. For panel **(D)** few of the contributing bacteria have been identified.

**Figure S5.**
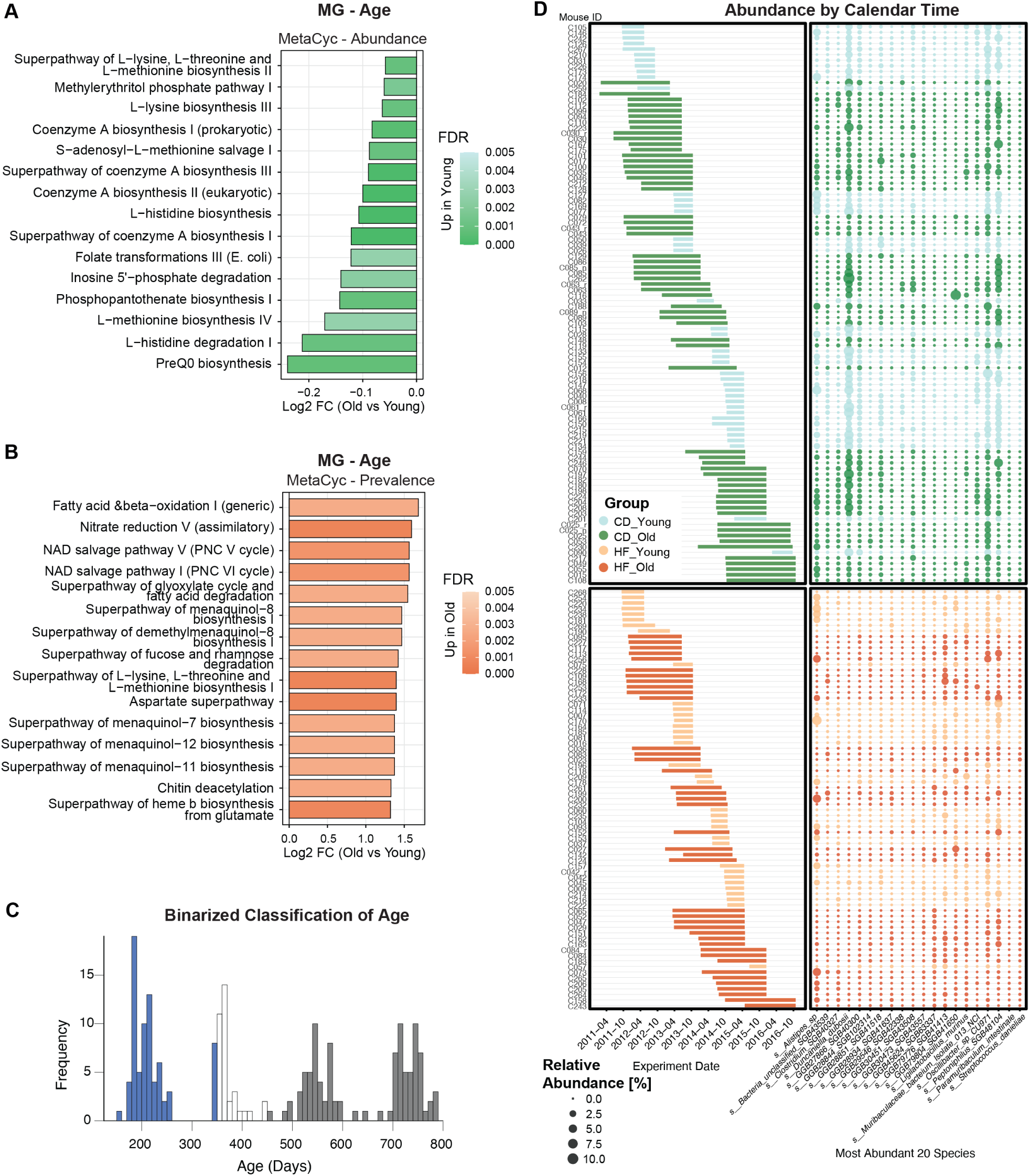
Supplementary Details Related to Age-Associated Analyses. (**A**) Abundance and (**B**) prevalence analysis of the top 15 age-associated MetaCyc pathways in MG data, showing fold changes and significances. (**C**) Distribution of sample ages across the MG cohort. Young (blue) and old (grey) groups were defined to ensure comparable sample sizes. Middle aged animals were not considered for the subsequent analysis on binarized age groups. (**D**) (Left) All mice (Y-axis) and the chronology of their timeline in the original aging colony, between 2011 and 2016, sorted by time and separated by dietary group (Right) The abundances of the top 20 species related to age (x-axis) across the same mice. No batch effect was observed due to time, i.e. there are no shifts in bacterial abundance according to the chronological years between 2011-2016.

**Figure S6.**
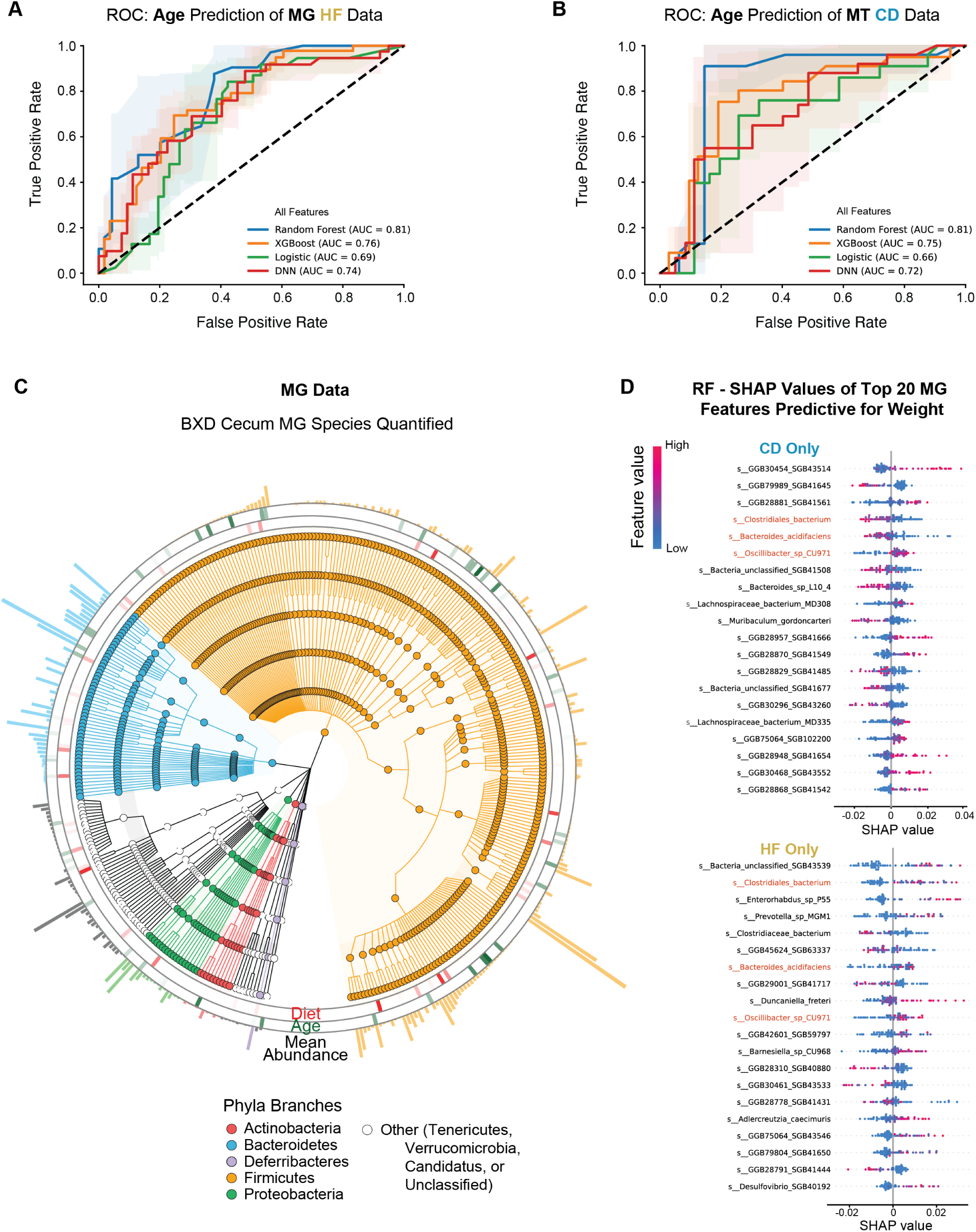
Machine Learning Models Predicting Age or Body Weight Using Multi-Omics Data. **(A)** ROC curves of different techniques to select features to develop models to predict age using the MG data for HF cohorts **(B)** and for CD cohorts. **(C)** Cladogram of the MG from the phylum (innermost ring of the dendogram, with 6 categories) to species (outermost ring of the dendogram) taxonomic rank. The innermost ring with bars represents species with significant differences across different diets. The second ring with bars represents species with significant differences across different ages. The third final ring indicates the mean relative abundance of each species. **(D)** Top 20 biomarkers selected by RF for body weight prediction using MG data, within CD (top) or within HF (bottom) cohorts. Three species selected in both cohorts are highlighted in red.

**Figure S7.**
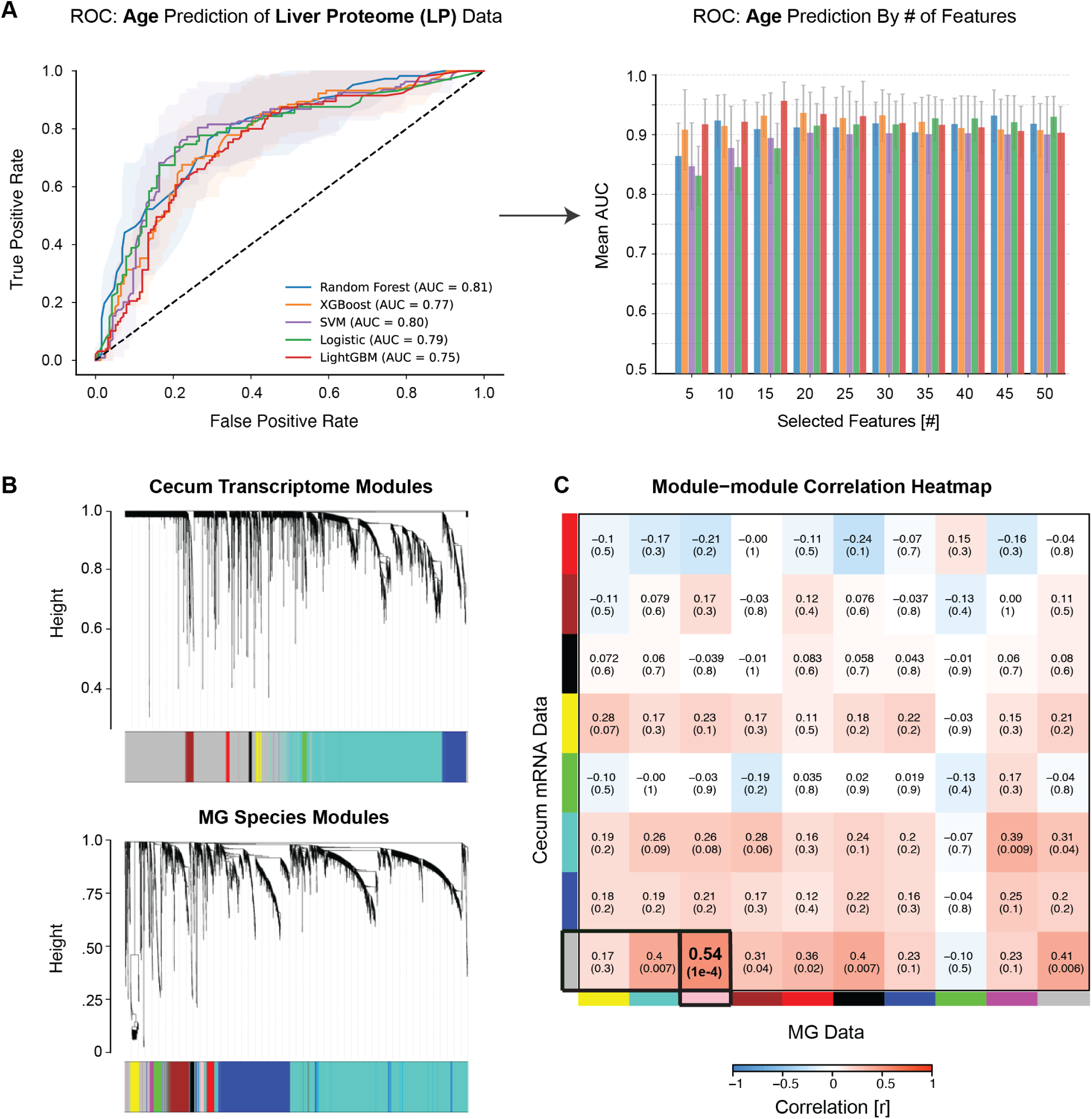
Supporting Results from Integrated Multi-omics and Network Analyses. **(A)** ROC curve for the various models for predicting age in CD mice (left) and bar plot for the numbers of features selected (right) using liver proteomics (LP) data. **(B)** Hierarchical clustering dendrograms of cecal mRNA (left panel) and MG (right panel), showing co-expression modules identified by WGCNA across all cecal transcripts (top) or MG species (bottom). **(C)** Module–module correlation heatmap between cecal mRNA modules and MG species modules. The strongest positive correlation was observed between MEgrey (cecum) and MEpink (microbiome) modules (r = 0.54, adj.p = 1e-4).

